# Comparative Analysis of Deep Learning Models for Predicting Causative Regulatory Variants

**DOI:** 10.1101/2025.05.19.654920

**Authors:** Gaetano Manzo, Kathryn Borkowski, Ivan Ovcharenko

## Abstract

**Motivation:** Genome-wide association studies (GWAS) have identified numerous noncoding variants associated with complex human diseases, disorders, and traits. However, resolving the uncertainty between GWAS association and causality remains a significant challenge. The small subset of noncoding GWAS variants with causative effects on gene regulatory elements can only be detected through accurate methods that assess the impact of DNA sequence variation on gene regulatory activity. Deep learning models, such as those based on Convolutional Neural Networks (CNNs) and transformers, have gained prominence in predicting the regulatory effects of genetic variants, particularly in enhancers, by learning patterns from genomic and epigenomic data. Despite their potential, selecting the most suitable model is hindered by the lack of standardized benchmarks, consistent training conditions, and performance evaluation criteria in existing reviews.

**Results:** This study evaluates state-of-the-art deep learning models for predicting the effects of genetic variants on enhancer activity using nine datasets stemming from MPRA, raQTL, and eQTL experiments, profiling the regulatory impact of 54,859 SNPs across four human cell lines. The results reveal that CNN models, such as TREDNet and SEI, consistently outperform other architectures in predicting the regulatory impact of single-nucleotide polymorphisms (SNPs). However, hybrid CNN-transformer models, such as Borzoi, display superior performance in identifying causal SNPs within a linkage disequilibrium block. While fine-tuning enhances the performance of transformer-based models, it remains insufficient to surpass CNN and hybrid models when evaluated under optimized conditions.

## Introduction

Genome-wide association studies (GWAS) have revealed that around 95% of disease-associated genetic variants occur in non-coding regions of the human genome, with causative variants commonly affecting regulatory elements that modulate gene expression [1, 2, 3]. These regulatory variants can profoundly affect phenotypes and alter disease susceptibility by dysregulating their target genes [4].

Deep learning has emerged as a transformative approach for predicting the regulatory effects of genetic variants, particularly within enhancer regions. These models harness large-scale genomic and epigenomic datasets to learn complex sequence-to-function relationships, identifying DNA sequence features that influence regulatory activity. Convolutional neural networks (CNNs), for example, have been successfully applied to detect variants that disrupt transcription factor binding sites or alter chromatin accessibility, offering mechanistic insights into their potential phenotypic consequences. Notable CNN-based models include DeepSEA [5], SEI [6], and TREDNet [7]. More recently, transformer-based architectures have demonstrated strong performance in capturing long-range dependencies and modeling cell-type-specific regulatory effects. These models—such as the DNABERT series [8, 9], the Nucleotide Transformer family [10, 11], and Enformer [12]—were pre-trained on large-scale genomic sequences using self-supervised objectives and subsequently fine-tuned for specialized tasks, including predicting DNA methylation patterns, enhancer activity, and the functional impact of disease-associated variants. By integrating contextual information across broader genomic regions, transformer models offer enhanced resolution for interpreting non-coding variation in a cell-type-aware manner. Selecting the most suitable model for detecting the regulatory effects of genetic variants remains a significant challenge despite several surveys offering detailed overviews of the deep learning ecosystem in this domain [13, 14, 15]. These surveys have highlighted the unique strengths and limitations of various models. However, they lack a unified framework for assessment. Specifically, existing reviews often fail to benchmark models on a standardized dataset, train or fine-tune them under consistent conditions, and evaluate their performance using uniform criteria. Furthermore, there is a fundamental difference between benchmarking models on regulatory regions versus regulatory variants, which is rarely addressed. While regulatory region analyses focus on identifying broader functional elements, regulatory variant assessments require evaluating the impact of specific sequence alterations within these regions, presenting distinct challenges and opportunities for model evaluation.

In this study, we evaluate state-of-the-art deep learning models to predict the effects of genetic variants on enhancer activity in the human genome. Our approach involved curating and integrating nine datasets derived from diverse experimental methodologies, including massively parallel reporter assay (MPRA), reporter assay quantitative trait loci (raQTL), and expression quantitative trait loci (eQTL) studies. These datasets encompass 54,859 single-nucleotide polymorphisms (SNPs) in enhancer regions across four human cell lines.

We tackle three tasks: predicting log2-fold changes in enhancer activity, classifying SNPs based on their regulatory impact, and identifying causal SNPs within linkage disequilibrium (LD) blocks. We also assess how data variability affects model performance, emphasizing the importance of data quality. Finally, we illustrate how enhancer detection models can be effectively leveraged for variant effect assessment, highlighting their practical utility in interpreting genomic data and prioritizing candidate regulatory variants.

To ensure a robust and thorough assessment, we fine-tuned 22 deep learning models for each cell line, systematically exploring a broad spectrum of architectural designs and parameter configurations. Our results indicate that CNN models outperform more “advanced” architectures, such as transformers, on causative regulatory variant detection. However, fine-tuning significantly boosts the performance of transformer-based architectures, revealing their potential to surpass CNNs under optimized conditions.

## Results

### Comparison of Enhancer Variant Prediction Models

We evaluated a diverse set of state-of-the-art deep learning models for their ability to predict the effects of genetic variants on enhancer activity in the human genome. These models differ markedly in architecture, parameter count, and the types of data used during training or pre-training. Notably, they were originally developed with distinct objectives in mind. For instance, models like Nucleotide Transformer [10] and DNABERT2 [9] were designed for broad genomic representation learning, whereas others—such as Geneformer [13] and TREDNet [7]—were tailored for more specific applications.

Geneformer, in particular, is a transformer-based model initially built to analyze single-cell gene expression profiles. However, it can be fine-tuned for a range of tasks, including chromatin state prediction and *in silico* perturbation modeling [13]. In our study, we adapted and fine-tuned Geneformer and other models to predict the effects of regulatory variants, leveraging their capacity to capture biological context for accurate assessment of enhancer activity changes (see *Materials and Methods*, Model Adaptation and Fine-tuning).

Importantly, this work compares not only model architectures but also their training foundations—highlighting how differences in pre-training data and objectives can impact downstream performance. To better disentangle these factors, we additionally assessed how the same model performs when applied across different experimental datasets. Despite their original design differences, all selected models were repurposed to predict enhancer variant effects (see *Materials and Methods*).

Our benchmark spans 54,859 single-nucleotide polymorphisms located in enhancer regions, compiled from nine datasets across four human cell lines: K562, HepG2, NPC, and HeLa. This comprehensive evaluation provides insights into how both model design and training data influence predictive accuracy for regulatory variant interpretation.

We performed regression analyses between model predictions and experimental log₂- fold changes for variant effects in enhancers across the human genome sequence. For each dataset, we conducted a separate analysis and averaged the results to provide a robust comparison across experimental settings (Pearson Correlation Fig. 1, Spearman Correlation Fig. S1, Materials and Methods). We acknowledge that our classification groups together all models that are not purely CNN- or transformer-based, despite significant architectural differences, particularly in the case of HyenaDNA, which diverges substantially from attention-based models. Nonetheless, our primary objective is to establish a broad guideline for enhancer variant prediction models, rather than to define rigid model categories.

**Fig 1:**
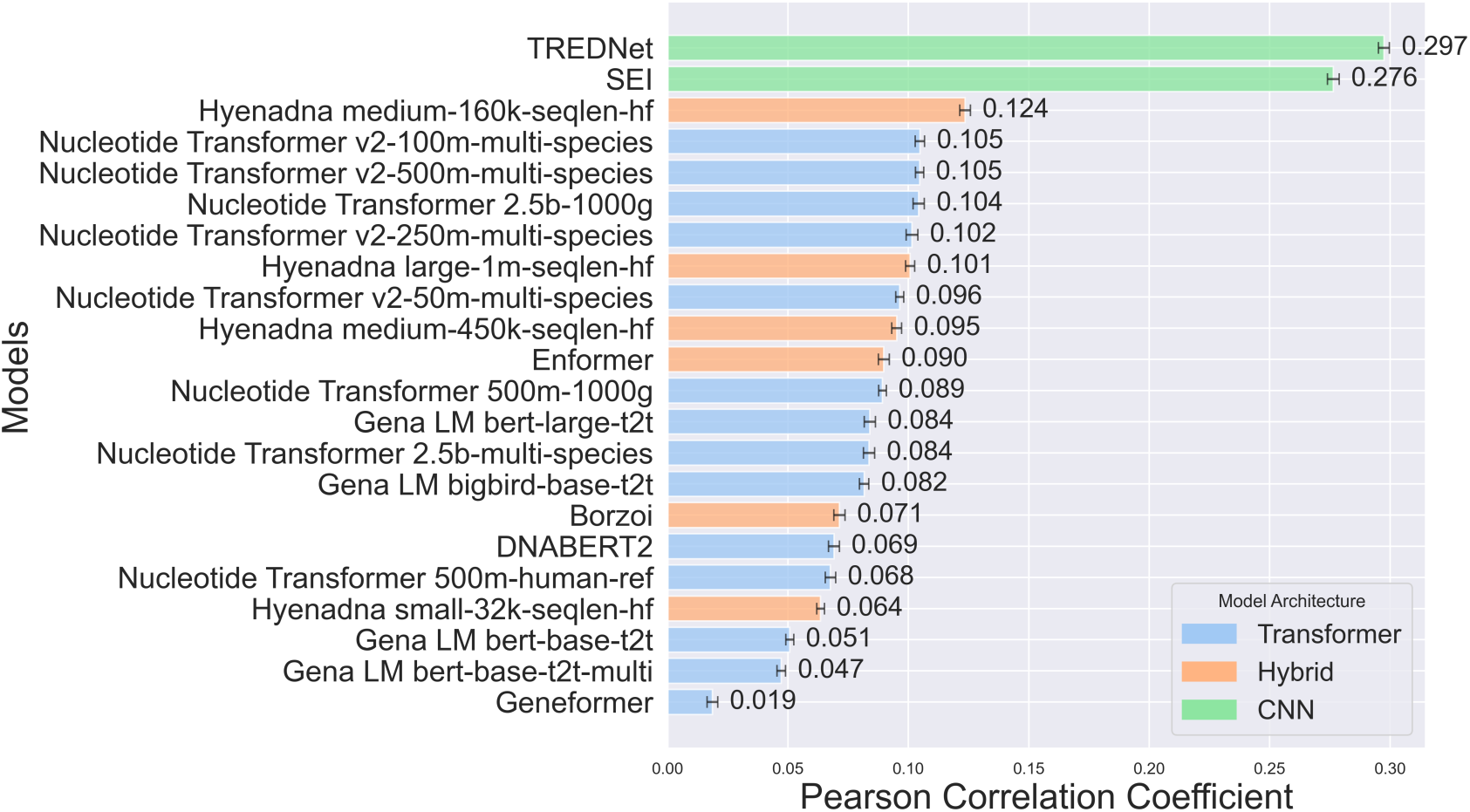
Pearson correlation coefficients between model predictions and experimental log2-fold difference for enhancer variant effects across the human genome. Bar colors denote model architectures (CNN: green, transformer: blue, and hybrid: orange). All correlations have p-values < 0.05, and error bars show variance.

CNN-based models, such as TREDNet and SEI, demonstrate the best alignment with experimentally recorded variant effects, achieving Pearson correlation coefficients of 0.297 and 0.276, respectively, while the most accurate transformer-based model, Nucleotide Transformer v2, scores only at 0.105, well below the level of statistical uncertainty. These findings highlight the effectiveness of CNNs in recognizing spatial and structural genomic patterns essential for predicting enhancer variant effects.

Convolutional layers identify genomic features, such as local motifs and epigenetic markers, that influence genetic variant effects [16]. This capability is crucial for capturing the complex genomic relationships underlying gene regulation and enhancer activity.

In contrast, transformer-based models fine-tuned for specific cell lines show lower correlation values (0.019–0.105). Despite their ability to model long-range relationships and complex sequences, they struggle to detect the subtle genomic patterns linked to regulatory variant effects.

This challenge likely stems from the complexity of genomic data, lower granularity in feature detection compared to CNNs, and the extensive data required to fully exploit transformer capabilities.

To bridge the gap between local pattern recognition and long-range dependency modeling, hybrid models combine components such as CNNs, transformers, and LSTMs. Models like the HyenaDNA series and Borzoi show moderate correlations, with HyenaDNA medium-160k-seqlen-hf achieving the highest among hybrids at 0.124. The integration of diverse components enhances performance compared to transformer-based models while optimizing computational costs relative to CNN-based models. However, despite their architectural versatility, these hybrid models do not surpass the performance of CNN-based models, highlighting a trade-off between flexibility and focused efficiency (Fig. 1, Fig. S1).

The classification task provides a deeper understanding of model performance by evaluating true positive and true negative rates in predicting variants that upregulate or downregulate gene expression (Fig. 2). Among 54,859 experimental SNPs, approximately half are linked to upregulation and half to downregulation.

**Fig 2:**
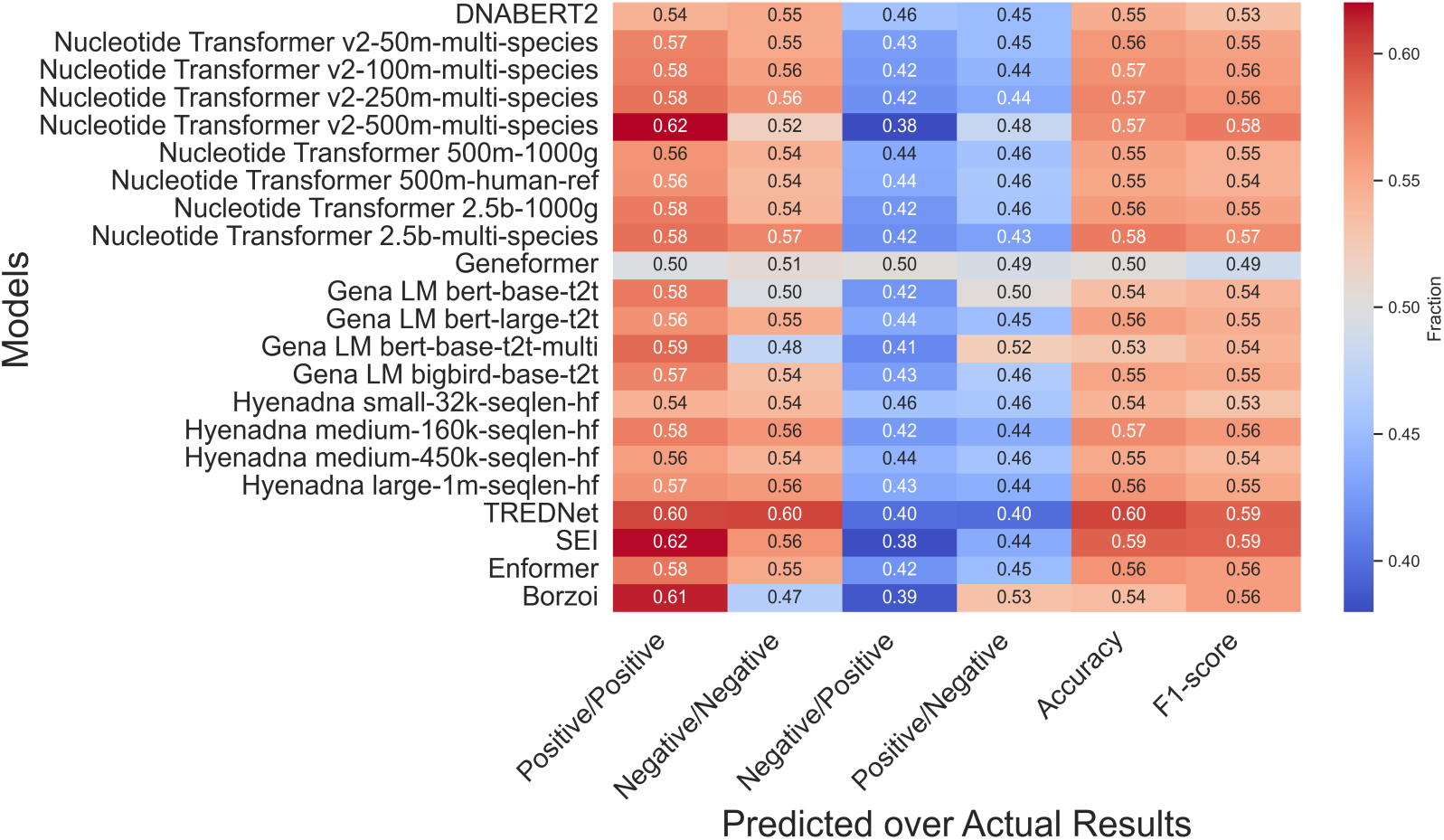
Heatmap of model variant predictions (Predicted) versus experimental values (Actual Results). The color intensity represents the fraction of values predicted as positive (negative) relative to the experimental positive (negative) values. Red indicates higher fractions (desired in the first two columns from the left), while blue indicates lower fractions (desired in the follow two columns). Accuracy and F1-score are indicated in the last two columns.

CNN-based models, like TREDNet and SEI, excel in predicting variant effects. TREDNet achieves balanced performance with correctly predicted upregulation (0.60) and downregulation (0.60) rates, while SEI leads in upregulation detection (0.62) but struggles with downregulation (0.56). Transformer-based models show mixed results. Nucleotide Transformer v2-500m-multi-species matches SEI’s upregulation prediction rate (0.62) but underperforms in downregulation (0.52). DNABERT2 performs steadily (0.54, 0.55), while Geneformer shows the weakest outcomes. Borzoi achieves a strong upregulation prediction rate (0.61) but falters in downregulation (0.47). Enformer shows moderate, balanced results (0.58, 0.55), while HyenaDNA remains consistent but unremarkable.

A crucial aspect of each model is its ability to identify causal SNPs within their respective Linkage Disequilibrium (LD) blocks. Essentially, this is the most important function of variant classification models, which can be directly applied to the resolution of causal variants within large blocks of associated variants identified in GWAS studies. To investigate this, we curated a dataset of 14,183 causal SNPs specific to the HepG2 cell line (Dataset 2, see Materials and Methods). For each causal SNP, we identified associated LD-block SNPs with an r^2^-value above 0.8 within a 500 kbp window, resulting in 263,286 SNPs. With this set, we can directly assess each method’s ability to retrieve the causal SNP within this ∼26:1 pool of associated SNPs. For each identified SNP, we generated two sequences of 1 kbp: one containing the SNP and the other containing the reference nucleotide. In both cases, the SNP or reference nucleotides were positioned at the center of the sequence. These sequences were then input into several models to assess their ability to predict causal SNPs by calculating the log2 fold change between the scores of the alternative sequence (containing the SNP) and the reference sequence (Methods and Materials sections). This procedure was repeated for all LD-block associated SNPs and the log2 fold-change score of the known causal SNP was compared to its associated variants. Finally, we computed the percentage of causal SNPs within an LD-block correctly predicted as causal based on their score being either the highest (“top-1” test) or within the top two and three highest scores (“top-2” and “top-3” tests).

The Borzoi model achieved the highest accuracy, correctly identifying 42.5% of causal SNPs, followed by the SEI, Enformer, and TREDNet models, with 40.7%, 39.4%, and 33.8% in the top1 test, respectively, (Fig. 3). The causal variant detection rate increased substantially in the top-2 and top-3 test with Borzoi (44.9%, 48.9), SEI (43.6%, 47.0%), Enformer (42.4%, 46.2%), and TREDNet (36.1%, 40.1%). Among the transformer-based models, the Gena LM bert-based-t2t-multi demonstrated the best performance (top-1 30.4%, top-2 32.7%, and top-3 37.5%), marginally surpassing other Nucleotide Transformer and Gena LM variants, which achieved accuracies between 28.0% and 30.0% in top-1 test. The HyenaDNA models, categorized as hybrid architectures, displayed moderate performance, with accuracies ranging from 24.3% to 26.5% in top- 1 test, and larger sequence lengths, such as those used in the HyenaDNA large model, led to improved results. The Geneformer model, however, had the lowest accuracy, identifying 24.3% of causal SNPs in top-1 test. Overall, hybrid and CNN architectures outperformed transformer-based models in this application, underscoring their suitability for tasks involving causal SNP detection.

**Fig 3:**
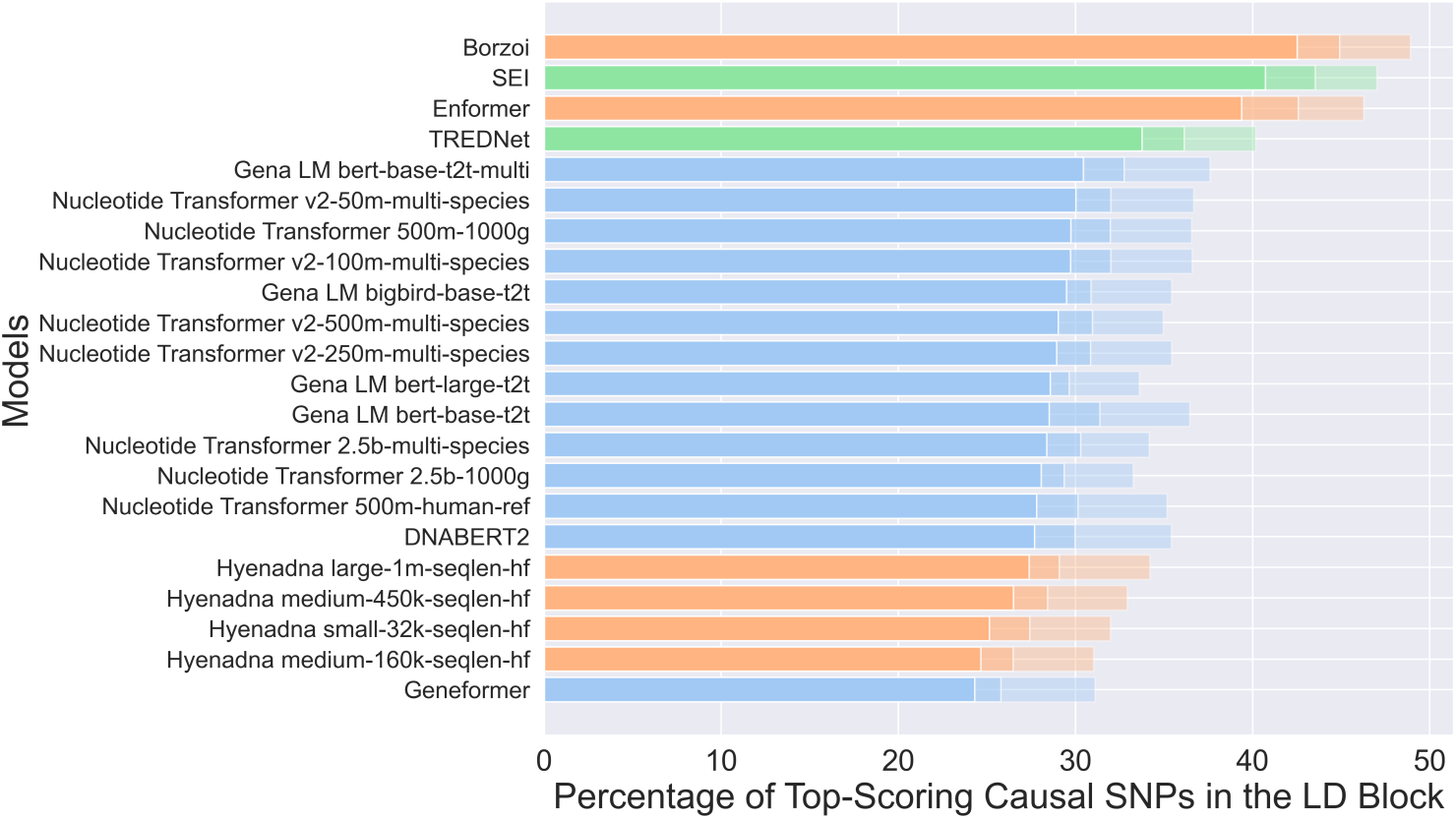
Percentage of top-scoring causal SNPs predicted within the Linkage Disequilibrium (LD) block by various models. Models are grouped by architecture: CNN (green), transformer (blue), and hybrid (orange). Bars are sorted by top-1 performance, with decreasing transparency indicating lower ranks (top-2 and top-3 results are progressively faded compared to top-1).

Overall, models like TREDNet and SEI consistently outperform others in both regression analysis and classification tasks, showcasing their strength in capturing local genomic patterns critical for regulatory variant effect prediction. Hybrid models, such as Borzoi and Enformer, offer balanced performance, bridging the gap between local pattern recognition and long-range dependency modeling but fail to surpass CNNs in overall accuracy. Yet, these models excel at accurate detection of causal variants within LD-blocks of associated SNPs. Transformer-based models, despite their potential to model complex sequences and long-range interactions, struggle to match the granularity of CNNs and hybrids.

### Impact of Certainty in Experimental Results

The results of experimental assays of regulatory variants depend on the selected degree of certainty in separating significant and insignificant variant effects. To investigate the impact of experimental data significance on correlation with modeling results, we analyzed the top models for each architecture type: TREDNet (CNN), HyenaDNA (hybrid), and Nucleotide Transformer (transformer). The experimental data was binned using the following significance thresholds, p < 0.5, p < 0.1, p < 5•10^-5^, p <10^-5^.

The results demonstrate that models align better with experimental findings when applied to variants with higher statistical significance. For instance, data points with greater experimental confidence show increased clustering in the first and third quadrants, representing cases where both experimental and predicted results are either positive (upregulation) or negative (downregulation) (Fig. 4, panels A-C, Figures S2, S3a, S3b, S3c, and S3d).

**Fig 4:**
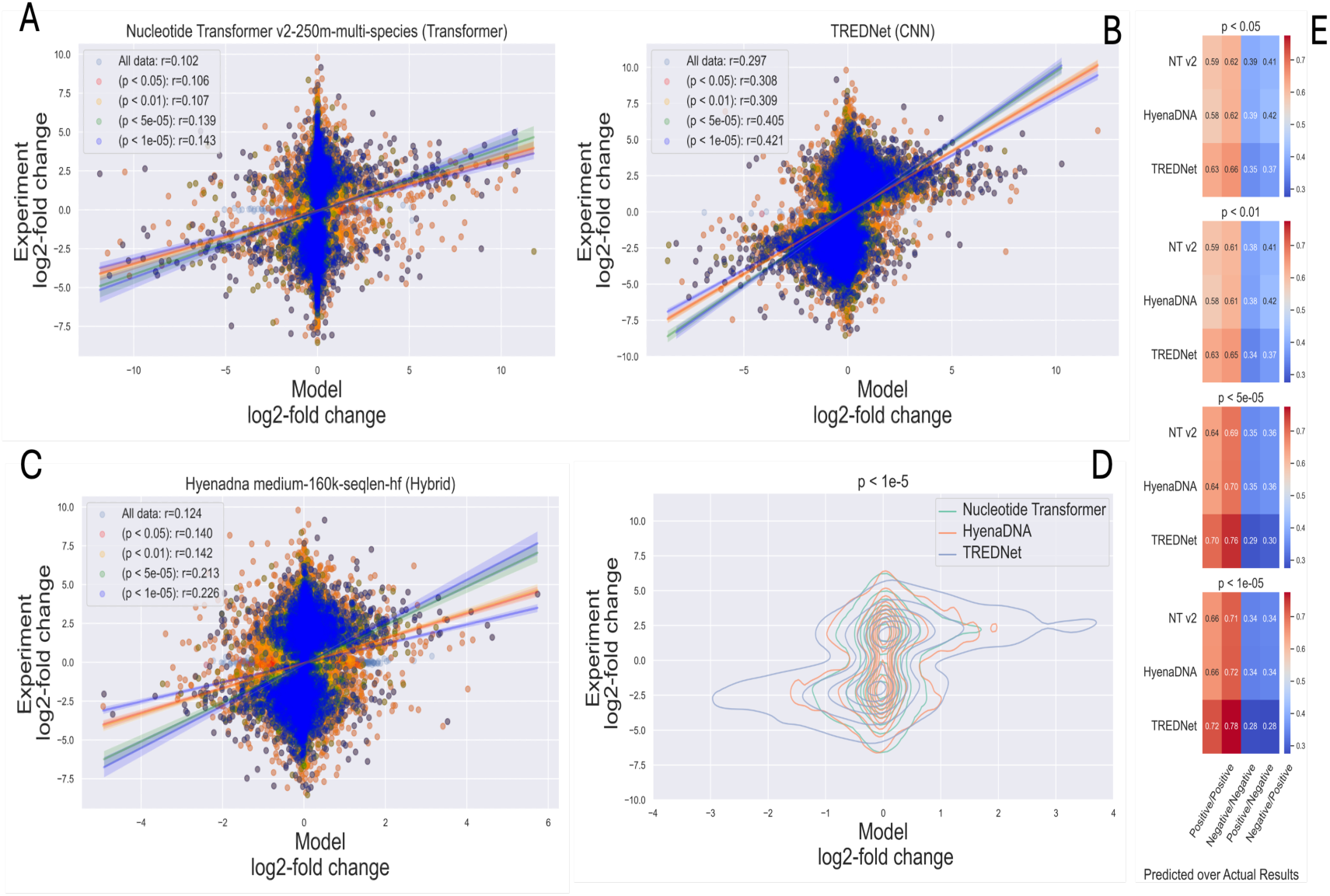
Model performance relative to experimental data significance, using top models from each architecture category. Panels A-C show scatter plots for TREDNet, HyenaDNA, and Nucleotide Transformer, with the correlation between model predictions and experimental results as significance improves. Panel D compares model predictions for highly significant variants. Panel E presents heatmaps of classification metrics at different p-value thresholds, highlighting performance variations in identifying positive/negative outcomes across architectures.

Among the evaluated architectures, HyenaDNA exhibited the greatest improvement in predictive accuracy with the increase of the statistical significance of experimental results (from all data 0.124 to 0.226 with p < 10^-5^ , Fig. 4, panel C). However, TREDNet consistently displayed superior overall performance, with a higher density of points aligning with experimental outcomes (from all data 0.298 to 0.421 with p < 10^-5^, Fig. 4, panel B). This trend is further illustrated in the density plots, where TREDNet exhibits increased clustering in the first and third quadrants, demonstrating stronger correlations between model predictions and experimental values compared to both hybrid and transformer models (Fig. 4, panel D). Moreover, the correlation coefficients across varying significance thresholds highlight TREDNet’s robustness, achieving the highest values under all conditions. True positives and true negatives increase with the significance of the experimental results (positive/positive 0.63, negative/negative 0.66, with p < 0.05; positive/positive 0.72, negative/negative 0.78, with p < 10^-5^), with the hybrid model and the transformer model following in performance (Fig. 4, panel E). The rest of models’ performance relative to experimental data significance can be found in Fig. S2, and Figures S3a, S3b, S3c, and S3d.

Given the strong cell-line specificity of enhancers (Wu and Huang, 2024), determining whether model performance depends on the type of cell line used is essential for understanding their predictive capabilities and limitations. For this, we evaluated whether a model fine-tuned for a specific cell line demonstrates superior performance compared to models from other architectures, such as CNNs, which often rely on generalization rather than cell-line-specific adaptation. Indeed, transformer-based models, pre-trained on extensive datasets and fine-tuned for targeted tasks, can offer both flexibility and accuracy, making them well-suited for capturing the unique regulatory landscape of cell lines. In contrast, CNN-based models, such as TREDNet and SEI, though highly effective for specific tasks, could require additional steps, such as retraining or the addition of specialized layers, to adapt their outputs for cell-line-specific predictions. However, the results indicate that TREDNet and SEI remain the most effective models for predicting variant effects across different cell lines, even after model fine-tuning (Pearson Correlation Table 1, Spearman Correlation Sup Table 1, Dataset-wise Sup Table 2 and Sup Table 3). For example, the Nucleotide Transformer v2-250m-multi-species model aligns with experimental results in K562 and HepG2 (0.1998 and 0.11) but performs less effectively in NPC and HeLa cells. This variability could stem from differences in pre-training datasets—such as multi-species versus human-specific data—and the fine-tuning approach.

**Table 1:** Pearson correlation coefficient for various deep learning models across four cell lines: K562 (19321 SNPs), HepG2 (16255 SNPs), NPC (14042 SNPs), and HeLa (5241 SNPs). Standard Error is reported in brackets. Bold and underline styles denote the top and second-highest correlations per cell line, respectively.

| Models | Cell Lines |  |  |  |
| --- | --- | --- | --- | --- |
|  | K562 | HepG2 | NPC | Hela |
|  | (19321 SNPs) | (16255 SNPs) | (14042 SNPs) | (5241 SNPs) |
| DNABERT2 | 0.086 (0.039) | 0.096 (0.191) | -0.004 (0.007) | 0.120 (0.140) |
| NT v2-50m-ms | 0.147 (0.058) | 0.103 (0.036) | 0.020 (0.003) | 0.074 (0.047) |
| NT v2-100m-ms | 0.152 (0.065) | 0.127 (0.050) | 0.016 (0.003) | 0.042 (0.027) |
| NT v2-250m-ms | 0.166 (0.064) | 0.110 (0.020) | 0.023 (0.003) | 0.030 (0.018) |
| NT v2 500m-ms | 0.199 (0.036) | 0.114 (0.007) | 0.028 (0.003) | 0.036 (0.011) |
| NT 500m1000g | 0.123 (0.084) | 0.116 (0.023) | 0.001 (0.002) | 0.022 (0.018) |
| NT 500m-h-ref | 0.149 (0.139) | 0.119 (0.038) | 0.013 (0.001) | 0.067 (0.043) |
| NT 2.5b-1000g | 0.147 (0.052) | 0.112 (0.021) | 0.005 (0.003) | 0.087 (0.105) |
| NT 2.5b-m-s | 0.153 (0.055) | 0.139 (0.027) | 0.027 (0.001) | 0.066 (0.027) |
| Geneformer | 0.005 (0.198) | 0.027 (0.160) | -0.002 (0.138) | -0.007 (0.057) |
| Gena LM-base | 0.077 (0.042) | 0.050 (0.033) | 0.024 (0.002) | 0.029 (0.227) |
| Gena LM large | 0.117 (0.044) | 0.088 (0.036) | 0.030 (0.002) | 0.069 (0.216) |
| Gena LM b-multi | 0.077 (0.051) | 0.062 (0.024) | 0.010 (0.001) | 0.026 (0.061) |
| Gena LM bigbird | 0.136 (0.055) | 0.099 (0.029) | 0.021 (0.002) | 0.045 (0.015) |
| Hyenadna 32k | 0.084 (0.071) | 0.098 (0.061) | 0.002 (0.002) | 0.031 (0.013) |
| Hyenadna 160k | 0.148 (0.053) | 0.142 (0.040) | -0.007 (0.003) | 0.046 (0.018) |
| Hyenadna 450k | 0.077 (0.074) | 0.134 (0.050) | -0.007 (0.003) | 0.055 (0.015) |
| Hyenadna 1mf | 0.118 (0.056) | 0.136 (0.059) | 0.001 (0.003) | 0.046 (0.013) |
| TREDNet | 0.315 (0.025) | 0.314 (0.018) | 0.075 (0.003) | 0.076 (0.063) |
| SEI | 0.297 (0.022) | 0.298 (0.029) | 0.073 (0.003) | 0.280 (0.008) |
| Enformer | 0.059 (0.059) | 0.141 (0.110) | 0.008 (0.080) | 0.245 (0.027) |
| Borzoï | 0.033 (0.016) | 0.125 (0.025) | 0.023 (0.004) | -0.066 (0.005) |

**Table 2:** Adaptation strategies for each deep learning model used in enhancer classification and variant effect prediction tasks. The table outlines how each model was adapted or applied to handle 1 kb genomic sequences, including preprocessing steps, tokenizer usage, fine-tuning status, and inference approaches. References correspond to the original sources describing each model.

| Models | Adaptation Strategy |
| --- | --- |
| DNABERT 2 [9] | Directly fine-tuned using Hugging Face's 'Trainer' on 1 kb genomic sequences using the pre-trained BPE tokenizer. Model input length was restricted to 512 tokens. Adapted |
|  | for binary classification (enhancer vs. control) across biosamples. Its ALiBi positional encoding and Flash Attention enabled efficient fine-tuning without modification to input format. |
| Nucleotide Transformer [10, 11] | Adapted using Hugging Face's 'Trainer' with 6-mer tokenization on 1 kb enhancer/control sequences. The tokenizer processed sequences into 512 tokens. No architectural changes were required, and the model's multi-species pretraining proved robust for human sequence classification. |
| Geneformer [13] | Originally trained on single-cell transcriptomic data using gene-level inputs, Geneformer was adapted for genomic language modeling by tokenizing 1 kb enhancer/control sequences using the pretrained vocabulary and tokenizer. Tokens mapped sequence chunks to "gene-like" units, enabling use of the transformer for DNA sequence classification. Fine-tuning was performed using Hugging Face's 'Trainer', with input sequences truncated or padded to 512 tokens. Despite domain shift, the model generalized well. |
| GENA-LM [17] | Fine-tuned on 1 kb genomic sequences using its native sparse attention tokenizer (BigBird). Used Hugging Face-compatible tokenization pipelines for BERT-base, BERT-large, and BigBird variants. Hugging Face 'Trainer' enabled flexible training for binary classification. Input sequences were processed to fit within the model's max length. |
| Enformer [12] | As the model was designed for very long sequences (up to 200 kb), adaptation involved truncating input to 1 kb and padding as needed. Preprocessing followed the authors' GitHub pipeline with minimal changes. No fine-tuning was performed—predictions were extracted directly from the pre-trained model for enhancer/control sequences. |
| HyenaDNA [18] | Used pre-trained weights and inference pipeline from the official GitHub repository. Inputs were standardized to 1 kb and padded to reach 32,768 tokens, as required by the model's architecture. Model was used in inference-only mode to evaluate predictions without fine-tuning. |
| Borzoi [19] | Applied directly on 1 kb input sequences using the GitHub-published inference code. Adaptation involved formatting enhancer/control sequences to match input requirements (one-hot or token-based). The model was not fine-tuned but used as-is for classification via regression output interpretation. |
| SEI [6] | Used as a non-fine-tuned model. 1 kb input sequences were one-hot encoded and passed through the SEI inference pipeline, which includes dilated convolutional layers for multi-scale pattern detection. Outputs were extracted from the final regulatory activity scores across 40+ cell types. |
| TREDNet [7] | Adapted for enhancer classification and SNP prioritization using the published pipeline. The model was inference-only; sequences were formatted to one-hot encoding and input into the TREDNet CNN. Saturated mutagenesis was used to evaluate variant effects post-classification. |

**Table 3:**
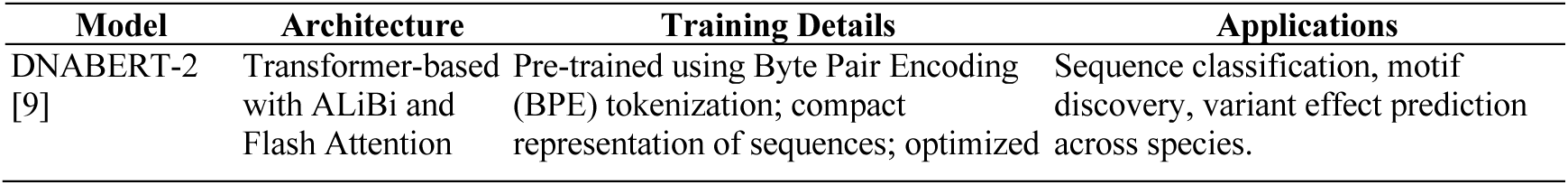
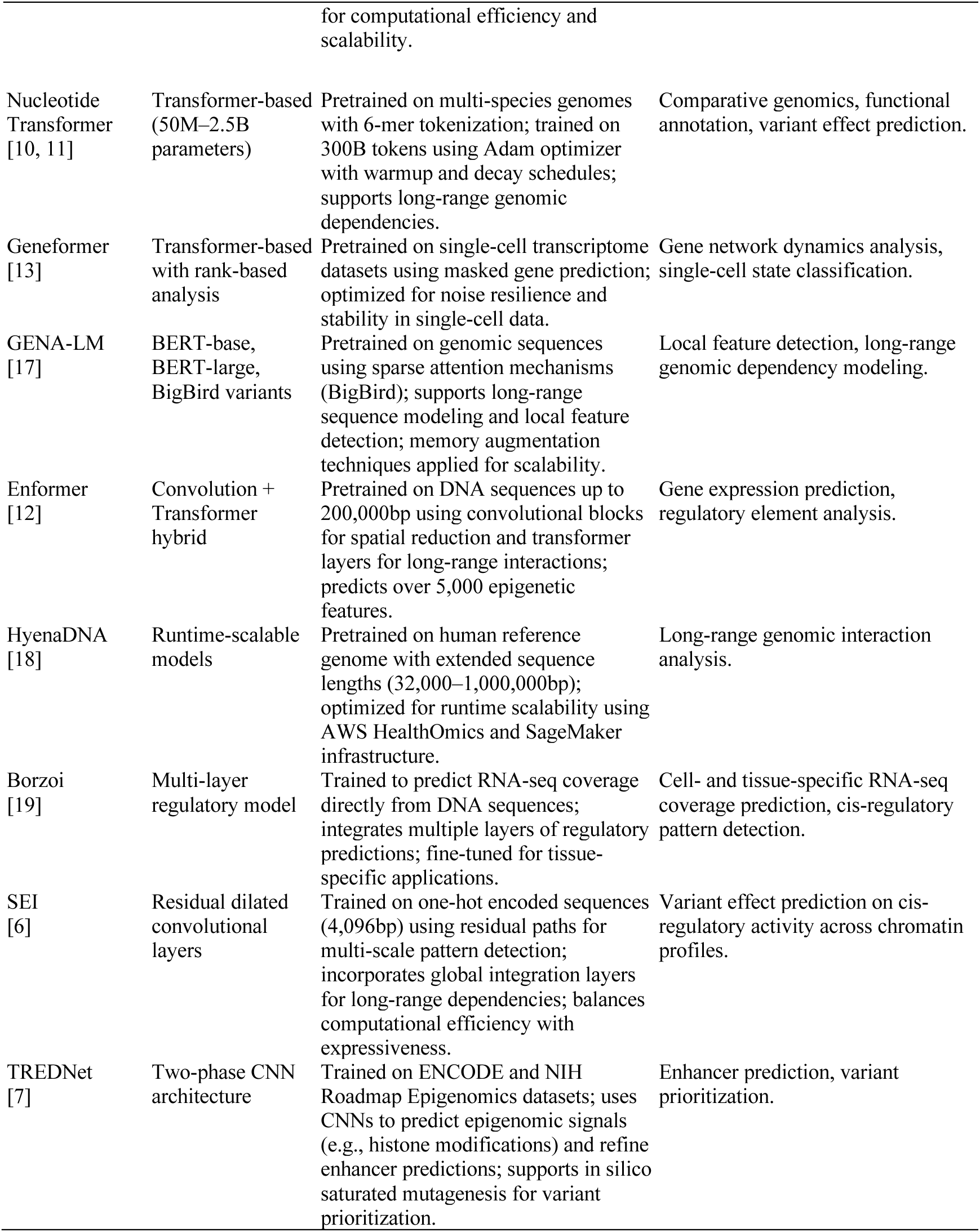
Models, architectures, training details, and applications of the models adopted.

Notably, the hybrid model Enformer, which integrates features from transformers and CNNs, performs particularly well in HeLa cells (0.245), ranking second only to SEI (0.279). As observed in the previous test, CNN-based models demonstrate superior performance, showcasing robustness across cell-line variants. We also assess the impact of the certainty of experimental results, SEI, which does not require training or fine-tuning for specific cell lines, outperforms fine-tuned models. TREDNet, likely benefiting from its second-phase training tailored to the specific cell line, achieves the best overall performance, further highlighting its ability to leverage cell-line-specific information effectively.

### Enhancer Detection Models for Variant Effect Assessment

Models designed to predict variant effects rely primarily on the change in confidence when detecting the enhancer region after introducing the variant. Next, we investigated whether improving the accuracy of enhancer sequence detection enhances predictions of variant effects. While SEI, Borzoi, and Enformer do not require fine-tuning, TREDNet, despite being a CNN-based model, incorporates a two-phase training process. The second phase can be considered fine-tuning [7].

Transformer-based architectures, particularly Nucleotide Transformer models pre-trained on multi-species datasets, emerge as strong performers in enhancer classification (Fig. S4). Models such as Nucleotide Transformer v2-250m-multi-species and Nucleotide Transformer 2.5b-multi-species consistently demonstrate high accuracy, as indicated by their auROC and auPRC metrics (auROC: 0.967, 0.963; auPRC: 0.958, 0.951). However, the correlation between the models’ outputs and the MPRA experimental results, which measure the fold-change effect between reference and alternative sequences, remains relatively low (ranging from 0.112 to 0.15). SEI stands out as the top performer in detecting enhancer activity, achieving the highest auROC (0.974) and auPRC (0.971) scores among all models. It is also the second best for the correlation between model outputs and experimental results (0.295). TREDNet demonstrates strong performance with a higher correlation (0.318), while balancing high auROC (0.872) and auPRC (0.814) scores. The HyenaDNA models show competitive classification metrics, with the Hyenadna medium-160k-seqlen-hf and Hyenadna large-1m-seqlen-hf versions attaining notable auROC (0.933, 0.904) and auPRC (0.921, 0.893) scores. However, their Pearson correlation coefficients are moderate (0.152, 0.151), indicating that while these models classify enhancer activity effectively, further optimization is needed to improve their predictive accuracy for variant-specific effects. Borzoi performs the weakest in detecting enhancer regions (auROC: 0.672, auPRC: 0.65), while Geneformer fares poorly in detecting variants (auROC: 0.027). Yet, this weak enhancer detection performance doesn’t translate in inaccurate detection of causal variants, as Borzoi outperforms all other methods in this critical test.

Fine-tuned models consistently outperform their one-shot counterparts, achieving F1 scores between 0.85 and 0.92, compared to 0.70 to 0.80 for one-shot models (Fig. 5, top). This improvement in performance is reflected in the correlation between model predictions and experimental results, particularly in detecting enhancer variants (Fig. 5, bottom). The performance enhancement is consistent across most architectures, with CNN-based models showing remarkable results even without fine-tuning. However, SEI, which is available only in its ’one-shot’ implementation, achieves a high F1 score and correlation, surpassing many fine-tuned transformer models. Additionally, some transformer-based models, such as Nucleotide Transformer 2.5b-1000g, show no improvement in detecting enhancer variants.

**Fig 5:**
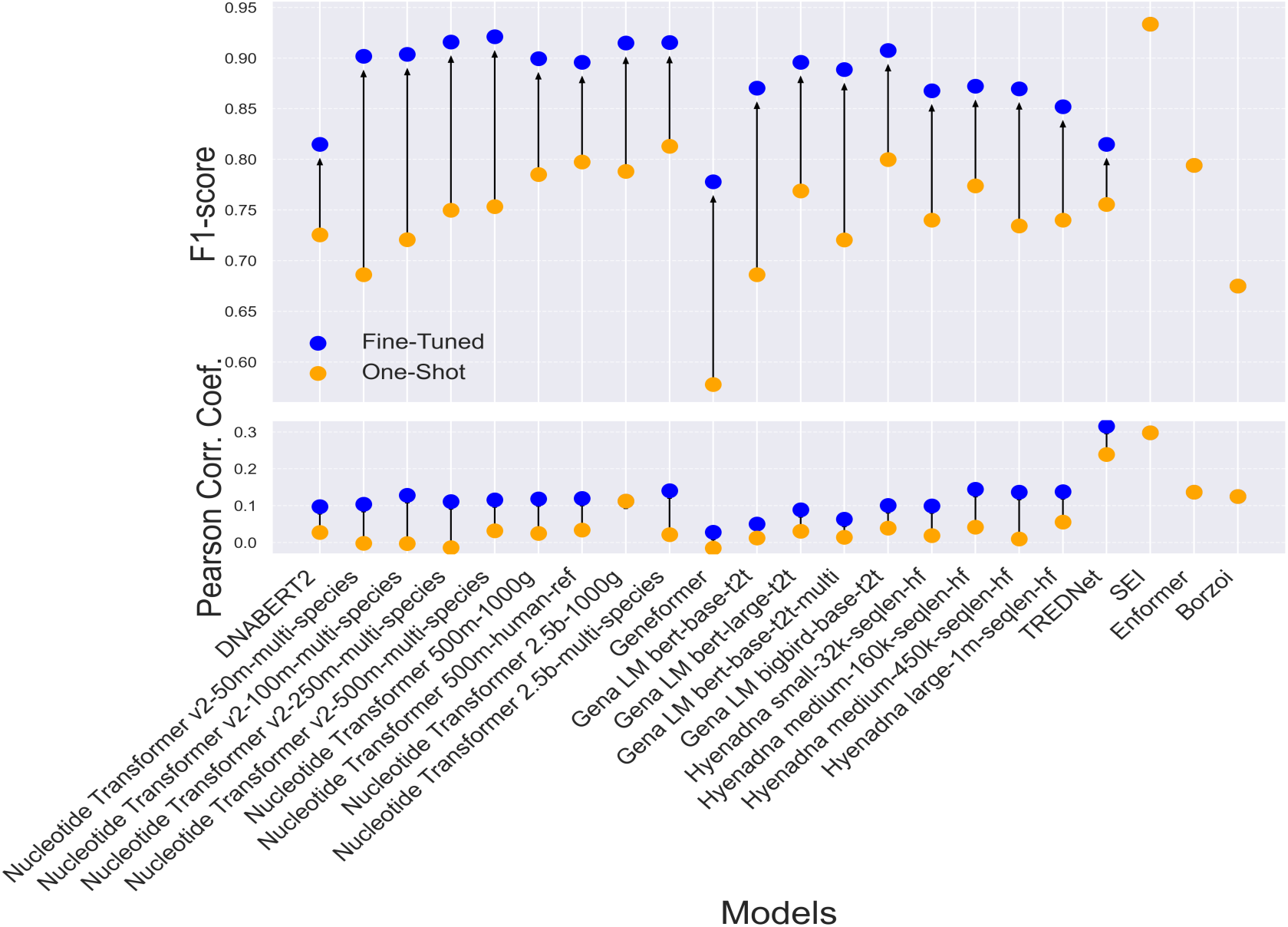
Comparison of F1 score (top) and Pearson Correlation Coefficient (bottom) between fine-tuned (blue dots) and one-shot (orange dots) models across various architectures, evaluated on Dataset 2, HepG2 cell line, (14183 SNPs). Arrows indicate the changes in performance between fine-tuned and one-shot implementations for each model.

## Discussion

Our evaluation of deep learning models highlights the strengths and limitations of different architectures in predicting the effects of genetic variants on enhancer activity. CNN-based models, such as TREDNet and SEI, outperform transformer-based and hybrid architectures in both regression and classification tasks. Notably, Borzoi, a hybrid model, excels in causal SNP prediction, leveraging both local and global genomic features critical for regulatory variant effect prediction.

### CNNs-, Transformers-, and Hybrid-based Models

TREDNet and SEI achieve superior performance in regression (Fig. 1) and classification tasks (Fig. 2). TREDNet demonstrates balanced predictive capability for both upregulated and downregulated variants, while SEI excels in detecting upregulated variants. CNNs effectively capture local genomic patterns like motifs and epigenetic markers, making them well-suited for enhancer activity analysis. Their simpler architecture yields high Pearson correlation coefficients (0.297 for TREDNet, 0.276 for SEI). However, their inability to model long-range dependencies limits performance in tasks requiring broader sequence context.

Transformers, including the Nucleotide Transformer and DNABERT 2, capture long-range dependencies but struggle with fine-grained regulatory signals. They underperform in enhancer variant effect prediction, with accuracies between 28.0% and 30.0% in causal SNP identification (Fig. 3). These models require substantial training data, and their performance could improve with expanded datasets or refined focus on regulatory signals.

Hybrid models, such as HyenaDNA and Borzoi, integrate CNNs, transformers, and LSTMs to balance local and long-range dependencies. While HyenaDNA shows moderate correlation with experimental results, Borzoi outperforms other models in causal SNP detection (Fig. 3), making it valuable for GWAS analysis. However, hybrid models do not consistently surpass CNNs in overall predictive accuracy and should be assessed based on task-specific requirements.

### Impact of Experimental Data Quality and Model Generalization

The reliability of model predictions is closely tied to the quality of experimental data. Model alignment with experimental results improves as statistical confidence in variant effects increases (Fig. 4). TREDNet, for instance, maintains high predictive accuracy even under varying significance thresholds, emphasizing the robustness of CNN-based models. These findings suggest that enhancing experimental rigor and statistical refinement can reduce uncertainty in variant effect assessments, leading to more reliable predictions.

In addition to data quality, model generalization across different cell lines is a critical factor. Transformer models fine-tuned for specific cell lines can capture regulatory landscapes but often underperform compared to CNNs. For instance, the Nucleotide Transformer v2-250m-multi-species achieves reasonable results in K562 and HepG2 but struggles in NPC and HeLa, likely due to pre-training limitations. In contrast, CNN models generalize well across cell lines without requiring fine-tuning, further demonstrating their robustness in variant effect prediction.

Enhancer detection models, such as the Nucleotide Transformer v2-250m-multi-species, achieve high classification accuracy (auROC: 0.967, auPRC: 0.958), yet their predictions show weak correlation with experimental results (Fig. S4). CNN-based models, particularly SEI, perform well in both enhancer detection and variant effect prediction. While fine-tuning benefits transformer models, SEI, despite its ‘one-shot’ implementation, still outperforms many fine-tuned transformers (Fig. 5). These results highlight the advantages of CNN architectures in both enhancer activity analysis and cross-cell-line generalization, reinforcing their suitability for regulatory variant effect prediction.

In conclusion, our findings suggest that SEI and TREDNet are the most effective models for predicting the effects of genetic variants on enhancer activity. Hybrid models like Borzoi demonstrate strong potential in extracting causal SNPs from GWAS data, showcasing their value in applications that require the integration of both local and global sequence contexts. Transformer-based models, on the other hand, are particularly effective at identifying enhancer regions due to their ability to capture long-range dependencies, although they still face challenges with fine-grained genomic features and often require substantial data for optimal performance. These results highlight the strengths of different model architectures for various aspects of regulatory variant prediction and underscore the need for further optimization to enhance their performance. Future work should aim to improve feature detection in transformer models and refine the flexibility of hybrid architectures to better capture both local and long-range genomic dependencies.

## Materials and Methods

### Data Pre-Processing

The datasets from HepG2, K562, NPC, and HeLa cell lines underwent a standardized preprocessing pipeline to ensure consistency and feature extraction for downstream analysis. Two main pipelines were applied: one for training and testing enhancer classifier models, such as Nucleotide Transformers and DNABERT, and another for mutagenesis experiments. For the enhancer classification pipeline, 1kb sequences were extracted for both enhancer (positive) and control regions.

To maintain consistency across models, all nucleotides were capitalized, as some models are case-sensitive. Unknown nucleotides marked as “N” were removed from the sequences. The datasets were then split into training, validation, and test sets, and tokenized using the corresponding model’s tokenizer with a maximum length of 512 tokens. Labels were generated to distinguish between positive and control sequences. In the mutagenesis experiment pipeline, the reference and alternative nucleotide locations were extracted for each row of the dataset. For each nucleotide of interest, a 1kb sequence was generated with the target nucleotide positioned in the center. Chromosome and position information, along with reference and alternative alleles, were also extracted. Log2 activity metrics, such as fold-change values or effect sizes, were calculated to assess regulatory activity. Additionally, signed p-values were derived from raw or adjusted p-values to capture both the statistical significance and directionality of the regulatory effects.

The preprocessing step ensured that variations in dataset structure did not interfere with downstream analysis. To further enhance consistency, hg19 genomic coordinates were used for variant mapping, and enhancer activity metrics were uniformly applied across datasets. Differences in expression or activity levels between experimental conditions, such as mutant versus control or modern versus archaic sequences, were computed when applicable. Ultimately, all datasets were converted into a standardized format that included chromosome information, positions, reference and alternative alleles, log2 activity ratios, signed p-values, raw p-values, and other relevant regulatory features.

### Model Adaptation and Fine-tuning

The objective of fine-tuning is to adapt models originally designed as enhancer classifiers to assess the impact of SNPs on enhancer activity. By introducing variant sequences, we evaluate how nucleotide changes affect model scores. To achieve this, we fine-tuned pre-trained DNA language models—including Geneformer (initially developed for scRNA-seq data)—as enhancer classifiers (Table 2).

Sequences were preprocessed as described earlier, with a length of 1 kbp, balancing genomic context and model compatibility. This length also reflects the average enhancer size of ∼400 bp. When adapting models like Enformer (196 kb input) and Borzoi (512 kb input), which do not accept arbitrary sequence lengths, we retained their full input size. The 1 kb variant-centered sequence was inserted into representative genomic contexts of the appropriate size, and predictions were averaged over multiple such contexts. Yet, for evaluation settings like Borzoi’s sliding window analysis where the method natively supports shorter sequences, we used 1 kb inputs directly.

Most models were loaded from the Hugging Face Hub using AutoTokenizer and AutoModelForSequenceClassification. Exceptions such as TREDNet, Enformer, SEI, and Borzoi were sourced from GitHub and required custom pipelines for preprocessing and inference.

Fine-tuning was performed using the Hugging Face transformers library for flexible training and evaluation. Models such as HyenaDNA, Geneformer, Gena-LM, DNABERT2, and Nucleotide Transformers were trained via the Trainer API on four biosample-specific datasets: HeLa (BioS2), neural progenitor cells (NPC, BioS45), HepG2 (BioS73), and K562 (BioS74). Sequences were cleaned to remove ambiguous bases (Ns) and split into training, validation, and test sets. Model-specific tokenizers processed sequences with a maximum input length of 512 tokens. For non-fine-tuned models like SEI and Enformer, the same 1 kbp sequence length was used with padding when appropriate.

Key hyperparameters included a batch size of 8, a learning rate of 1e-5, and a maximum of 200 epochs. Early stopping with a patience of 5 epochs was used to avoid overfitting, with progress logged every 500 steps. Fine-tuned models were saved in output directories named by model checkpoint and biosample ID and uploaded to the Hugging Face Hub for reproducibility. GPU memory was cleared after each session for efficiency. This workflow enabled the adaptation of general-purpose DNA language models for cell-type-specific regulatory prediction while ensuring consistency with non-fine-tuned models. Training was performed on NIH BioWulf node clusters with A100 GPUs, enabling efficient handling of large datasets (e.g., ∼130,000 sequences for positive and control enhancer regions), which completed in hours using 8 A100 GPUs.

### Validation and Post-Processing

During the fine-tuning process, the models were optimized to distinguish enhancer sequences from control sequences. In contrast, the validation process focused on evaluating the difference between the reference enhancer sequence and the alternative sequence, where the middle nucleotide was mutated. The difference in output scores between these two sequences, obtained from the fine-tuned model, allowed for the estimation of the variant’s effect on the enhancer. If the output scores for both sequences were identical, the variant had no effect. Conversely, a change in output scores indicated either up- or down-regulation.

The prediction process involved generating regulatory effect predictions for each dataset by calculating the log2 ratio of alternative to reference allele probabilities for each variant. This metric provided a standardized approach to assess the relative regulatory effects of alternative alleles compared to reference alleles. Since each fine-tuned model produced logits for each class (enhancer or not), predictions for both reference and alternative alleles were processed by first calculating margin logits, which represent the difference in predicted class probabilities. These logits were then transformed into probabilities using the sigmoid function. The log2 ratio of these probabilities was computed to quantify the regulatory differences between the two alleles. Additionally, precomputed predictions stored in pickle files were loaded for specific models and datasets. The predictions for reference and alternative alleles were used to compute a ratio, representing the alternative-to-reference prediction ratio. This ratio was subsequently log2-transformed to ensure consistency in the output format. Additional preprocessing steps were applied to specific datasets as needed, such as loading specialized prediction files.

The prediction pipeline was applied across all datasets by iterating through experiments defined in a metadata table. For each experiment, data were preprocessed, and predictions were either computed or loaded accordingly. The results were compiled into a unified dictionary containing log2 prediction ratios for all variants across experiments. As in the fine-tuning process, the validation of variant/reference sequences was performed using 1 kb sequences, ensuring consistency in genomic context and input length across models. Several metrics were employed to evaluate model performance, including the Area Under the Curve (AUC), Precision-Recall Curve (PRC), F1-score, and correlation coefficients such as Pearson and Spearman. These metrics provided a comprehensive assessment of model performance, encompassing both classification accuracy and the correlation between predicted and observed values.

### Deep Learning Models

This study utilizes several deep learning models designed for genomic sequence analysis, with a particular focus on applications to the Homo sapiens genome. The models represent the forefront of innovation in deep learning for genomics, leveraging diverse architectures and pre-training paradigms to handle the complexity of human genomic data. To maintain focus on models explicitly designed for the human genome, this study excluded models like EVO and GPN, which are primarily aimed at other species or biological domains.

### Training Data

We constructed a dataset of positive and control enhancer sequences from four human cell lines: HepG2, Hela, K562, and Neural Progenitor Cells (NPC). These cell lines were selected for their relevance in genomic research and the availability of comprehensive epigenomic data, such as DNase-Seq and histone modification profiles, sourced from the ENCODE project using genome annotations for hg19 and hg38.

Positive enhancer sequences were defined through a two-step process. First, open chromatin regions were identified using DNase-Seq data. These regions were then intersected with H3K27ac histone modification signals, a hallmark of active enhancers. To refine the dataset, we excluded non-enhancer regions such as exonic regions (coding sequences), promoter regions (near transcription start sites), and low-confidence regions listed in the ENCODE Blacklist (e.g., repetitive sequences). Specific blacklisted peaks from UCSC browser tables were also removed.

Control sequences were selected using a similar filtering process but specifically excluded regions overlapping both DNase-Seq peaks and H3K27ac signals to ensure they represented non-active genomic regions. Both positive and control sequences underwent identical exclusion criteria to eliminate confounding genomic features such as exons and promoters.

All sequences were standardized to 1 kb in length to balance sufficient coverage of regulatory regions with computational efficiency. This size is widely used in enhancer studies as it captures relevant regulatory elements while avoiding extraneous genomic features. The dataset includes an equal number of positive and control sequences for each cell line: HepG2 (14,062 each), NPC (12,466 each), K562 (19,968 each), and Hela (31,247 each). Metadata from the ENCODE project was used to retrieve biosample IDs and associated DNase-Seq and histone modification data. Reference files provided coordinates for exons, blacklist regions, and promoters, ensuring only valid enhancer or control regions were included in the final dataset.

### Validation Data

To systematically characterize the functional effects of genetic variants on gene regulation, we compiled a diverse set of high-throughput reporter assay datasets spanning multiple cell types, genomic contexts, and experimental designs (Table 4).

**Table 4:**
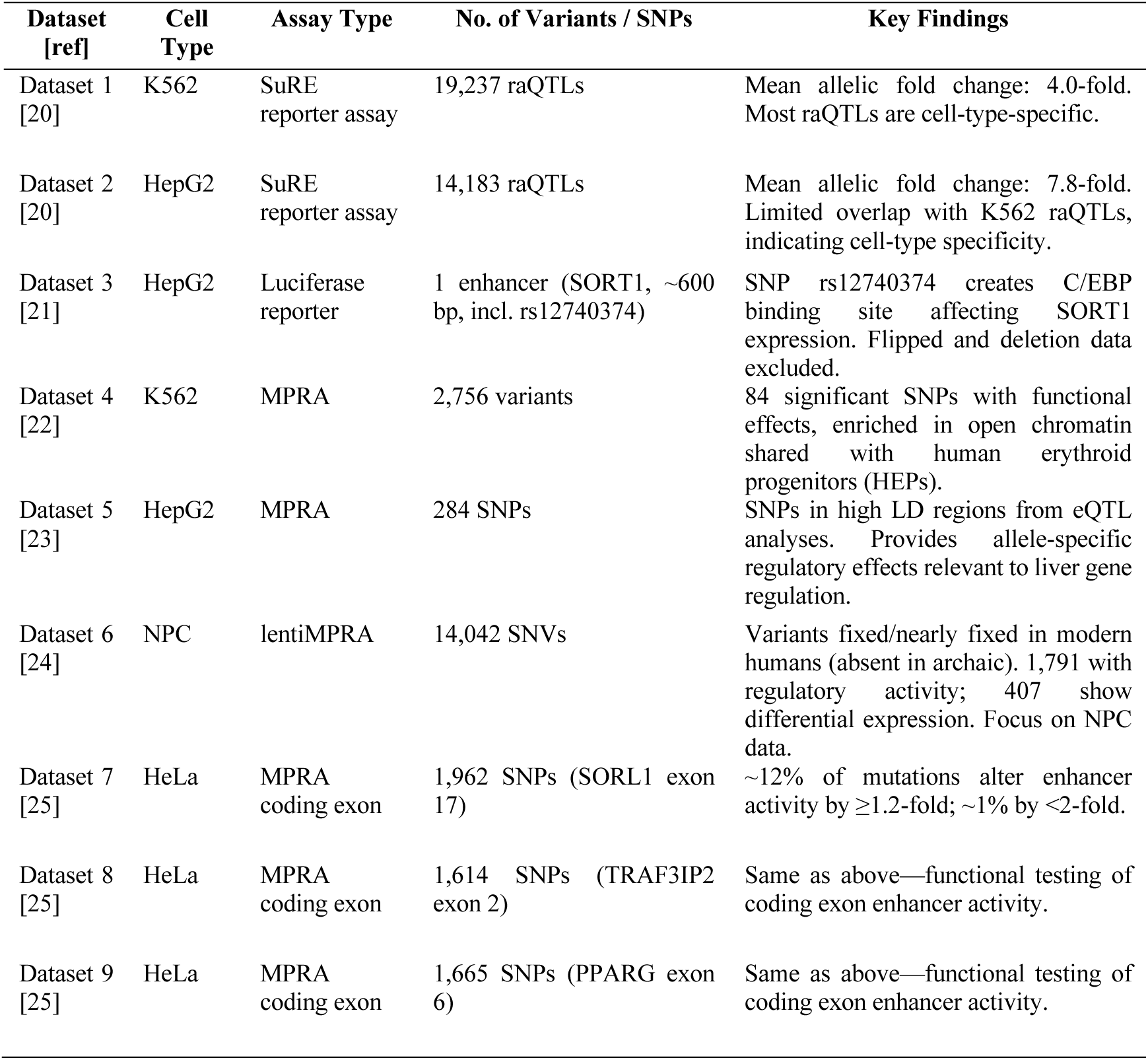
Overview of the datasets used to assess regulatory variant function across diverse cell types and genomic contexts.

Datasets 1 and 2 were sourced from the study by [20], which utilized the Survey of Regulatory Elements (SuRE) reporter assay to identify reporter assay quantitative trait loci (raQTLs)—SNPs influencing regulatory element activity. For K562 cells, 19,237 raQTLs were identified with an average allelic fold change of 4.0-fold, while HepG2 cells yielded 14,183 raQTLs with a higher average fold change of 7.8-fold. Most raQTLs were cell-type-specific, showing limited overlap between the two cell lines.

Dataset 3 was derived from [21] and focuses on a ∼600 bp enhancer region at the SORT1 locus (1p13). This region includes SNP rs12740374, which creates a C/EBP binding site that influences SORT1 expression. The enhancer was tested in HepG2 cells using a luciferase reporter assay. Data for flipped sequences were excluded due to the orientation-independent nature of enhancers, and deletion datasets were omitted due to computational model constraints, which focus on single-nucleotide variants.

Dataset 4, from [22] used MPRA to investigate 2,756 variants linked to red blood cell traits in K562 cells. The study identified 84 highly significant SNPs with functional effects, enriched in open chromatin regions shared between K562 cells and primary human erythroid progenitors (HEPs). This dataset provides insights into erythroid-specific regulatory activity.

Dataset 5, generated by [23], includes MPRA results for 284 SNPs tested in HepG2 cells. These SNPs, located within regions of high linkage disequilibrium identified through eQTL analyses, offer a detailed map of allele-specific regulatory effects relevant to liver-specific gene regulation and genotype-phenotype associations.

Dataset 6, from [24], includes lentiMPRA results for 14,042 single-nucleotide variants fixed or nearly fixed in modern humans but absent in archaic humans (e.g., Neanderthals). Tested across three cell types—embryonic stem cells, neural progenitor cells, and fetal osteoblasts—the study identified 1,791 sequences with regulatory activity, including 407 with differential expression between modern and archaic humans. This dataset focuses on NPC data.

Datasets 7, 8, and 9, from [25], include MPRA results for three coding exons—SORL1 exon 17, TRAF3IP2 exon 2, and PPARG exon 6—tested in HeLa cells. Respectively, these datasets cover 1,962, 1,614, and 1,665 SNPs. The study explored enhancer activity within these coding exons (eExons), finding that ∼12% of mutations altered enhancer activity by at least 1.2-fold, with ∼1% causing changes of less than 2-fold.

### Declarations

#### Ethics approval and consent to participate

Not applicable.

### Consent for publication

Not applicable.

### Availability of data and material

Code, data, and analysis repository: https://github.com/tanoManzo/AI4Genomic.git

Fine-tuned models: https://huggingface.co/tanoManzo

### Authors’ contributions

G.M. performed the computational analyses. K.B. curated and validated the experimental data, prepared figures, and contributed to data interpretation. I.O. conceptualized the study and established the experimental framework. All authors contributed to writing, editing, and revising the final manuscript. All authors reviewed the manuscript.

### Funding

This research was supported by the Division of Intramural Research of the National Library of Medicine, National Institutes of Health; 1-ZIA-LM200881-12 (to I.O.).

## Supporting information

Supporting Information - Comparative Analysis of Deep Learning Models for Predicting Causative Regulatory Variants

## Acknowledgements

This work utilized the computational resources of the NIH HPC Biowulf cluster (http://hpc.nih.gov).

## References

1. Uffelmann E, Huang QQ, Munung NS, de Vries J, Okada Y, Martin AR, et al. Genome-wide association studies. Nat Rev Methods Primers. 2021 Dec;1(1):59.

2. Visscher PM, Wray NR, Zhang Q, Sklar P, McCarthy MI, Brown MA, et al. 10 years of GWAS discovery: biology, function, and translation. Am J Hum Genet. 2017 Jul 6;101(1):5–22.

3. Knight JC. Approaches for establishing the function of regulatory genetic variants involved in disease. Genome Med. 2014 Oct 31;6(10):92.

4. Albert FW, Kruglyak L. The role of regulatory variation in complex traits and disease. Nat Rev Genet. 2015 Apr;16(4):197–212.

5. Zhou J, Troyanskaya OG. Predicting effects of noncoding variants with deep learning-based sequence model. Nat Methods. 2015 Oct;12(10):931–4.

6. Chen KM, Wong AK, Troyanskaya OG, Zhou J. A sequence-based global map of regulatory activity for deciphering human genetics. Nat Genet. 2022 Jul 11;54(7):940–9.

7. Hudaiberdiev S, Taylor DL, Song W, Narisu N, Bhuiyan RM, Taylor HJ, et al. Modeling islet enhancers using deep learning identifies candidate causal variants at loci associated with T2D and glycemic traits. Proc Natl Acad Sci USA. 2023 Aug 29;120(35):e2206612120.

8. Ji Y, Zhou Z, Liu H, Davuluri RV. DNABERT: pre-trained Bidirectional Encoder Representations from Transformers model for DNA-language in genome. Bioinformatics. 2021 Aug 9;37(15):2112–20.

9. Zhou Z, Ji Y, Li W, Dutta P, Davuluri R, Liu H. DNABERT-2: Efficient Foundation Model and Benchmark For Multi-Species Genome. arXiv. 2023;

10. Dalla-Torre H, Gonzalez L, Mendoza Revilla J, Lopez Carranza N, Henryk Grywaczewski A, Oteri F, et al. The nucleotide transformer: building and evaluating robust foundation models for human genomics. BioRxiv. 2023 Jan 15;

11. de Almeida BP, Dalla-Torre H, Richard G, Blum C, Hexemer L, Gelard M, et al. SegmentNT: annotating the genome at single-nucleotide resolution with DNA foundation models. BioRxiv. 2024 Mar 15;

12. Avsec Ž, Agarwal V, Visentin D, Ledsam JR, Grabska-Barwinska A, Taylor KR, et al. Effective gene expression prediction from sequence by integrating long-range interactions. Nat Methods. 2021 Oct 4;18(10):1196–203.

13. Theodoris CV, Xiao L, Chopra A, Chaffin MD, Al Sayed ZR, Hill MC, et al. Transfer learning enables predictions in network biology. Nature. 2023 Jun;618(7965):616–24.

14. Consens ME, Dufault C, Wainberg M, Forster D, Karimzadeh M, Goodarzi H, et al. To Transformers and Beyond: Large Language Models for the Genome. arXiv. 2023;

15. Alharbi WS, Rashid M. A review of deep learning applications in human genomics using next-generation sequencing data. Hum Genomics. 2022 Jul 25;16(1):26.

16. Yue T, Wang Y, Zhang L, Gu C, Xue H, Wang W, et al. Deep learning for genomics: from early neural nets to modern large language models. Int J Mol Sci. 2023 Nov 1;24(21).

17. Fishman V, Kuratov Y, Petrov M, Shmelev A, Shepelin D, Chekanov N, et al. GENA-LM: A Family of Open-Source Foundational Models for Long DNA Sequences. BioRxiv. 2023 Jun 13;

18. Nguyen E, Poli M, Faizi M, Thomas A, Birch-Sykes C, Wornow M, et al. HyenaDNA: Long-Range Genomic Sequence Modeling at Single Nucleotide Resolution. arXiv. 2023 Nov 14;

19. Linder J, Srivastava D, Yuan H, Agarwal V, Kelley DR. Predicting RNA-seq coverage from DNA sequence as a unifying model of gene regulation. BioRxiv. 2023 Sep 1;

20. van Arensbergen J, Pagie L, FitzPatrick VD, de Haas M, Baltissen MP, Comoglio F, et al. High-throughput identification of human SNPs affecting regulatory element activity. Nat Genet. 2019 Jul;51(7):1160–9.

21. Kircher M, Xiong C, Martin B, Schubach M, Inoue F, Bell RJA, et al. Saturation mutagenesis of twenty disease-associated regulatory elements at single base-pair resolution. Nat Commun. 2019 Aug 8;10(1):3583.

22. Ulirsch JC, Nandakumar SK, Wang L, Giani FC, Zhang X, Rogov P, et al. Systematic functional dissection of common genetic variation affecting red blood cell traits. Cell. 2016 Jun 2;165(6):1530–45.

23. Vockley CM, Guo C, Majoros WH, Nodzenski M, Scholtens DM, Hayes MG, et al. Massively parallel quantification of the regulatory effects of noncoding genetic variation in a human cohort. Genome Res. 2015 Aug;25(8):1206–14.

24. Weiss CV, Harshman L, Inoue F, Fraser HB, Petrov DA, Ahituv N, et al. The cis- regulatory effects of modern human-specific variants. eLife. 2021 Apr 22;10.

25. Birnbaum RY, Patwardhan RP, Kim MJ, Findlay GM, Martin B, Zhao J, et al. Systematic dissection of coding exons at single nucleotide resolution supports an additional role in cell-specific transcriptional regulation. PLoS Genet. 2014 Oct 23;10(10):e1004592.

