## Supporting Information - Comparative Analysis of Deep Learning Models for Predicting Causative Regulatory Variants for "Comparative Analysis of Deep Learning Models for Predicting Causative Regulatory Variants"

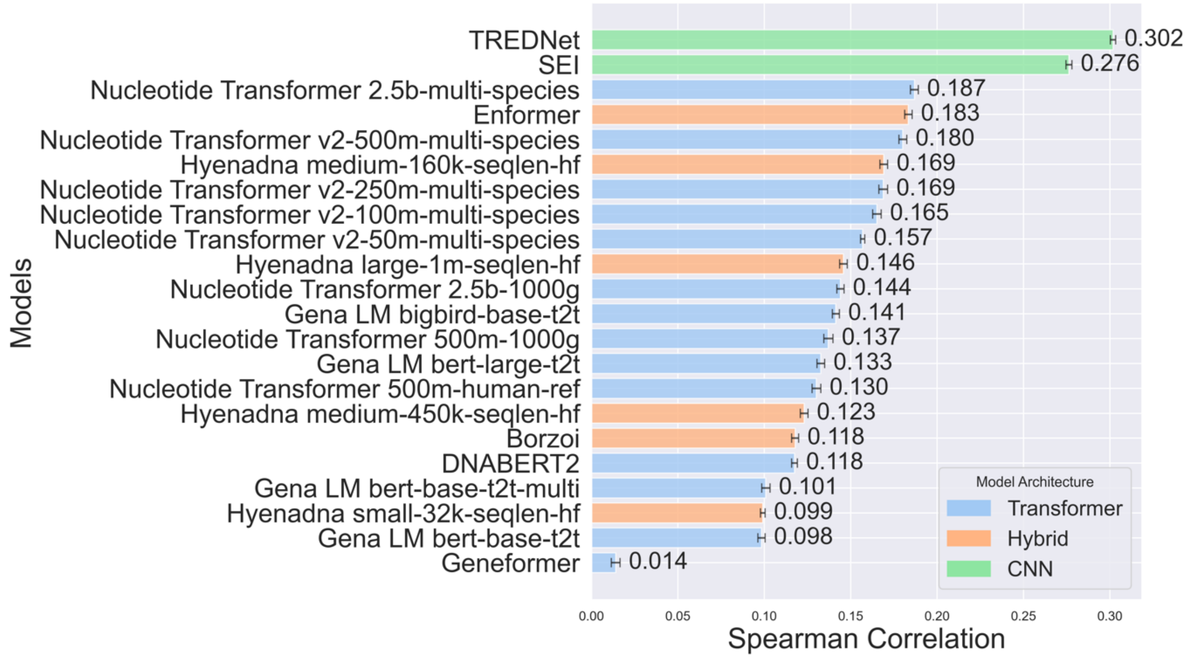


***Figure S1**. Spearman correlation between model predictions and experi-mental log2-fold changes for enhancer variant effects across the human genome. Bar colors denote model architectures (CNN: green, transformer: blue, and hybrid: orange). All correlations have p-values < 0.05, and error bars show variance.*

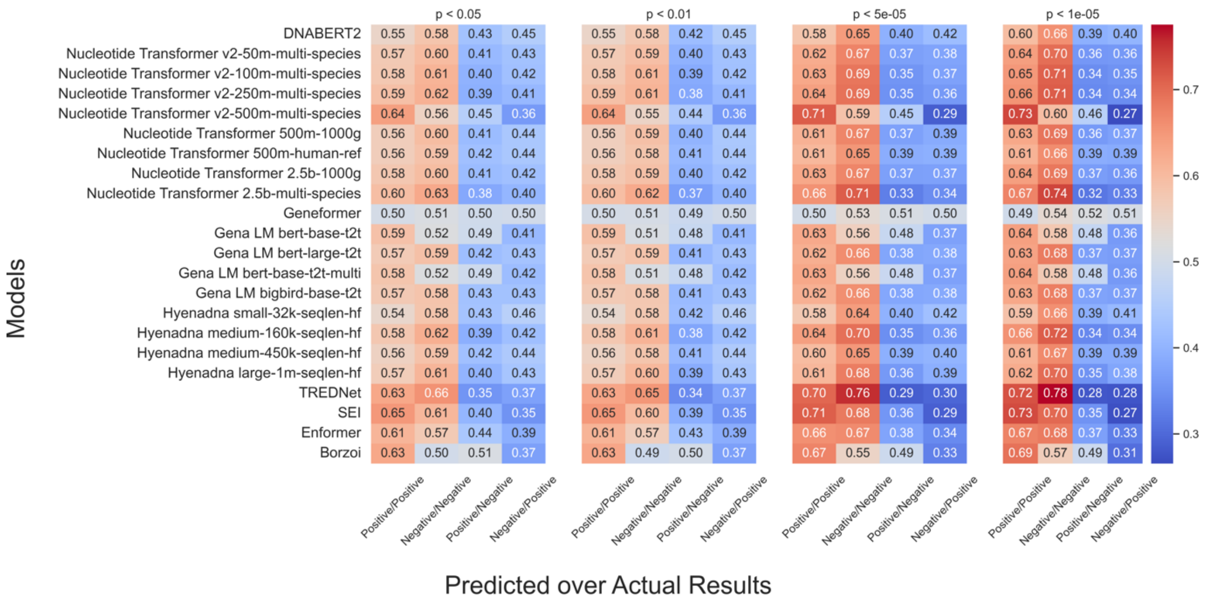


***Figure S2**. Heatmap of model variant predictions (Predicted) versus experimental values (Actual Results) at different p-value thresholds, highlighting performance variations in identifying positive/negative outcomes across architectures.. The color intensity represents the fraction of values predicted as positive/negative relative to the experimental positive/negative values. Red indicates higher fractions (desired in the first two columns from the left), while blue indicates lower fractions (desired in the last two columns).*

***Table S1.*** *Spearman correlation for various deep learning models across four cell lines: K562 (19321 SNPs), HepG2 (16255 SNPs), NPC (14042 SNPs), and HeLa (5241 SNPs). Bold and underline styles denote the top and second-highest correlations per cell line, respectively.*

| | **Cell Lines** | | | |
|---|---|---|---|---|
| **Models** | **k562 (19321 SNPs)** | **hepg2 (16255 SNPs)** | **NPC (14042 SNPs)** | **Hela (5241 SNPs)** |
| DNABERT2 | 0.136255 | 0.13844 | 0.00983 | 0.10131 |
| Nucleotide Transformer v2-50m-multi-species | 0.193606 | 0.176066 | 0.029745 | 0.091993 |
| Nucleotide Transformer v2-100m-multi-species | 0.194085 | 0.210449 | 0.029202 | 0.085566 |
| Nucleotide Transformer v2-250m-multi-species | 0.230211 | 0.178415 | 0.031789 | 0.091625 |
| Nucleotide Transformer v2-500m-multi-species | 0.255502 | 0.193536 | 0.039783 | 0.076763 |
| Nucleotide Transformer 500m-1000g | 0.152783 | 0.193222 | 0.010707 | 0.06466 |
| Nucleotide Transformer 500m-human-ref | 0.158163 | 0.17474 | 0.016139 | 0.058858 |
| Nucleotide Transformer 2.5b-1000g | 0.167759 | 0.181192 | 0.014801 | 0.081246 |
| Nucleotide Transformer 2.5b-multi-species | 0.226941 | 0.22058 | 0.016179 | 0.091921 |
| Geneformer | 0.00589 | 0.030644 | -0.00983 | -0.009303 |
| Gena LM bert-base-t2t | 0.122552 | 0.116882 | 0.025676 | 0.052523 |
| Gena LM bert-large-t2t | 0.169897 | 0.142243 | 0.041452 | 0.084151 |
| Gena LM bert-base-t2t-multi | 0.096464 | 0.140658 | 0.015713 | 0.047429 |
| Gena LM bigbird-base-t2t | 0.169682 | 0.159723 | 0.023648 | 0.088436 |
| Hyenadna small-32k-seqlen-hf | 0.110639 | 0.129993 | 0.000771 | 0.075399 |
| Hyenadna medium-160k-seqlen-hf | 0.1927 | 0.208351 | 0.007788 | 0.118122 |
| Hyenadna medium-450k-seqlen-hf | 0.115686 | 0.186035 | 0.007832 | 0.093602 |
| Hyenadna large-1m-seqlen-hf | 0.162212 | 0.194539 | 0.012185 | 0.077734 |
| TREDNet | **0.313784** | **0.365124** | **0.068044** | 0.103394 |
| SEI | <u>0.287206</u> | <u>0.315009</u> | <u>0.067235</u> | **0.155081** |
| Enformer | 0.131957 | 0.230396 | -0.005137 | <u>0.129005</u> |
| Borzoi | 0.089077 | 0.173757 | 0.024613 | -0.019793 |

***Table S2.*** *Pearson correlation coefficients for various deep learning models across multiple datasets (Data 1-9 and the respective SNPs). Bold and underline styles denote the top and second-highest correlations per cell line, respectively.*

| Model | Dataset 1 | Dataset 2 | Dataset 3 | Dataset 4 | Dataset 5 | Dataset 6 | Dataset 7 | Dataset 8 | Dataset 9 |
|---|---|---|---|---|---|---|---|---|---|
| | (19237 SNPs) | (14183 SNPs) | (1789 SNPs) | (84 SNPs) | (283 SNPs) | (14042 SNPs) | (1692 SNPs) | (1614 SNPs) | (1665 SNPs) |
| | K562 | HepG2 | HepG2 | G562 | HepG2 | NPC | Hela | Hela | Hela |
| DNABERT2 | 0.086 | 0.098 | 0.049 | 0.140 | 0.137 | -0.004 | 0.613 | 0.480 | 0.211 |
| Nucleotide Transformer v2-50m-multi-species | 0.147 | 0.104 | 0.129 | 0.178 | 0.322 | 0.021 | 0.575 | 0.354 | 0.218 |
| Nucleotide Transformer v2-100m-multi-species | 0.152 | 0.128 | 0.113 | 0.207 | **0.570** | 0.022 | 0.526 | 0.232 | 0.136 |
| Nucleotide Transformer v2-250m-multi-species | 0.166 | 0.111 | 0.217 | 0.142 | 0.431 | 0.042 | -0.383 | 0.306 | **0.329** |
| Nucleotide Transformer v2-500m-multi-species | 0.199 | 0.116 | -0.030 | 0.389 | 0.028 | 0.084 | 0.261 | 0.264 | -0.010 |
| Nucleotide Transformer 500m-1000g | 0.123 | 0.119 | 0.111 | 0.325 | 0.068 | 0.000 | 0.532 | 0.346 | 0.162 |
| Nucleotide Transformer 500m-human-ref | 0.149 | 0.120 | 0.052 | 0.296 | 0.427 | 0.002 | 0.449 | 0.253 | 0.218 |
| Nucleotide Transformer 2.5b-1000g | 0.147 | 0.113 | 0.165 | 0.238 | 0.182 | 0.004 | 0.607 | 0.357 | 0.188 |
| Nucleotide Transformer 2.5b-multi-species | 0.153 | 0.140 | 0.340 | 0.311 | 0.491 | 0.060 | 0.557 | 0.366 | 0.066 |
| Geneformer | 0.005 | 0.028 | 0.029 | 0.228 | 0.121 | 0.004 | -0.108 | -0.021 | 0.031 |
| Gena LM bert-base-t2t | 0.077 | 0.050 | -0.008 | 0.151 | -0.138 | 0.047 | -0.465 | 0.216 | 0.031 |
| Gena LM bert-large-t2t | 0.117 | 0.089 | 0.192 | 0.203 | 0.234 | 0.041 | 0.754 | 0.099 | 0.017 |
| Gena LM bert-base-t2t-multi | 0.076 | 0.063 | -0.058 | 0.156 | -0.081 | 0.004 | -0.400 | 0.057 | 0.107 |
| Gena LM bigbird-base-t2t | 0.136 | 0.100 | 0.291 | 0.412 | -0.013 | 0.039 | 0.624 | 0.338 | 0.017 |
| Hyenadna small-32k-seqlen-hf | 0.084 | 0.100 | 0.104 | 0.174 | 0.265 | 0.019 | 0.293 | 0.015 | 0.251 |
| Hyenadna medium-160k-seqlen-hf | 0.149 | 0.144 | 0.267 | 0.289 | 0.123 | -0.016 | 0.317 | 0.137 | 0.318 |
| Hyenadna medium-450k-seqlen-hf | 0.077 | 0.137 | 0.073 | 0.209 | 0.214 | -0.036 | 0.630 | 0.166 | 0.238 |
| Hyenadna large-1m-seqlen-hf | 0.118 | 0.138 | 0.126 | 0.243 | 0.204 | -0.016 | 0.606 | 0.116 | 0.227 |
| TREDNet | **0.316** | **0.342** | 0.397 | **0.601** | 0.363 | 0.167 | 0.784 | 0.410 | 0.085 |
| SEI | 0.298 | 0.298 | 0.394 | 0.341 | 0.051 | **0.190** | 0.801 | **0.595** | 0.237 |
| Enformer | 0.058 | 0.137 | **0.492** | 0.410 | 0.434 | 0.037 | **0.843** | 0.542 | 0.142 |
| Borzoi | 0.033 | 0.125 | 0.012 | 0.306 | 0.121 | 0.050 | 0.231 | -0.237 | -0.120 |

***Table S3.*** *Spearman correlation for various deep learning models across multiple datasets (Data 1-9 and the respective SNPs). Bold and underline styles denote the top and second-highest correlations per cell line, respectively.*

| Model | **Dataset 1** (19237 SNPs) **K562** | **Dataset 2** (14183 SNPs) **HepG2** | **Dataset 3** (1789 SNPs) **HepG2** | **Dataset 4** (84 SNPs) **G562** | **Dataset 5** (283 SNPs) **HepG2** | **Dataset 6** (14042 SNPs) **NPC** | **Dataset 7** (1692 SNPs) **Hela** | **Dataset 8** (1614 SNPs) **Hela** | **Dataset 9** (1665 SNPs) **Hela** |
|---|---|---|---|---|---|---|---|---|---|
| DNABERT2 | 0.136 | 0.136 | 0.103 | 0.168 | 0.105 | 0.018 | 0.627 | 0.513 | 0.336 |
| Nucleotide Transformer v2-50m-multi-species | 0.194 | 0.170 | 0.191 | 0.348 | -0.018 | 0.029 | 0.273 | 0.496 | 0.252 |
| Nucleotide Transformer v2-100m-multi-species | 0.194 | 0.203 | 0.222 | 0.135 | <u>0.382</u> | 0.016 | 0.545 | 0.442 | 0.069 |
| Nucleotide Transformer v2-250m-multi-species | 0.231 | 0.171 | 0.306 | 0.196 | 0.372 | 0.027 | 0.309 | 0.507 | 0.186 |
| Nucleotide Transformer v2-500m-multi-species | 0.256 | 0.187 | -0.020 | 0.442 | 0.226 | 0.079 | 0.264 | 0.432 | -0.089 |
| Nucleotide Transformer 500m-1000g | 0.153 | 0.188 | 0.148 | 0.429 | -0.077 | 0.014 | 0.345 | 0.390 | 0.064 |
| Nucleotide Transformer 500m-human-ref | 0.158 | 0.170 | 0.062 | <u>0.528</u> | 0.066 | 0.007 | 0.309 | 0.416 | 0.111 |
| Nucleotide Transformer 2.5b-1000g | 0.168 | 0.173 | 0.285 | 0.488 | 0.355 | 0.016 | 0.582 | 0.495 | 0.104 |
| Nucleotide Transformer 2.5b-multi-species | 0.228 | 0.209 | 0.342 | 0.400 | 0.305 | 0.040 | 0.427 | 0.517 | 0.072 |
| Geneformer | 0.006 | 0.030 | 0.044 | 0.330 | -0.076 | 0.007 | 0.027 | -0.039 | -0.017 |
| Gena LM bert-base-t2t | 0.122 | 0.115 | 0.066 | 0.285 | 0.180 | 0.059 | -0.518 | 0.394 | 0.091 |
| Gena LM bert-large-t2t | 0.170 | 0.135 | 0.221 | 0.510 | 0.360 | 0.047 | 0.555 | 0.448 | 0.199 |
| Gena LM bert-base-t2t-multi | 0.096 | 0.140 | 0.012 | 0.410 | -0.138 | 0.019 | -0.035 | 0.242 | 0.157 |
| Gena LM bigbird-base-t2t | 0.170 | 0.155 | 0.287 | 0.512 | 0.090 | 0.037 | 0.482 | 0.428 | 0.178 |
| Hyenadna small-32k-seqlen-hf | 0.111 | 0.126 | 0.200 | 0.391 | 0.146 | 0.011 | 0.545 | -0.096 | <u>0.371</u> |
| Hyenadna medium-160k-seqlen-hf | 0.193 | 0.201 | 0.310 | 0.487 | 0.178 | -0.009 | 0.409 | 0.461 | **0.428** |
| Hyenadna medium-450k-seqlen-hf | 0.116 | 0.181 | 0.147 | 0.274 | 0.167 | -0.013 | 0.655 | 0.376 | 0.326 |
| Hyenadna large-1m-seqlen-hf | 0.162 | 0.188 | 0.182 | 0.453 | 0.270 | -0.010 | <u>0.700</u> | 0.406 | 0.370 |
| TREDNet | **0.314** | **0.360** | 0.416 | **0.573** | **0.435** | <u>0.135</u> | 0.673 | 0.536 | 0.267 |
| SEI | <u>0.287</u> | <u>0.309</u> | <u>0.427</u> | 0.495 | 0.149 | **0.148** | 0.627 | **0.633** | 0.248 |
| Enformer | 0.132 | 0.218 | **0.566** | 0.374 | 0.267 | -0.016 | **0.900** | <u>0.553</u> | 0.053 |
| Borzoi | 0.088 | 0.172 | 0.036 | 0.270 | 0.173 | 0.073 | 0.282 | -0.368 | 0.009 |

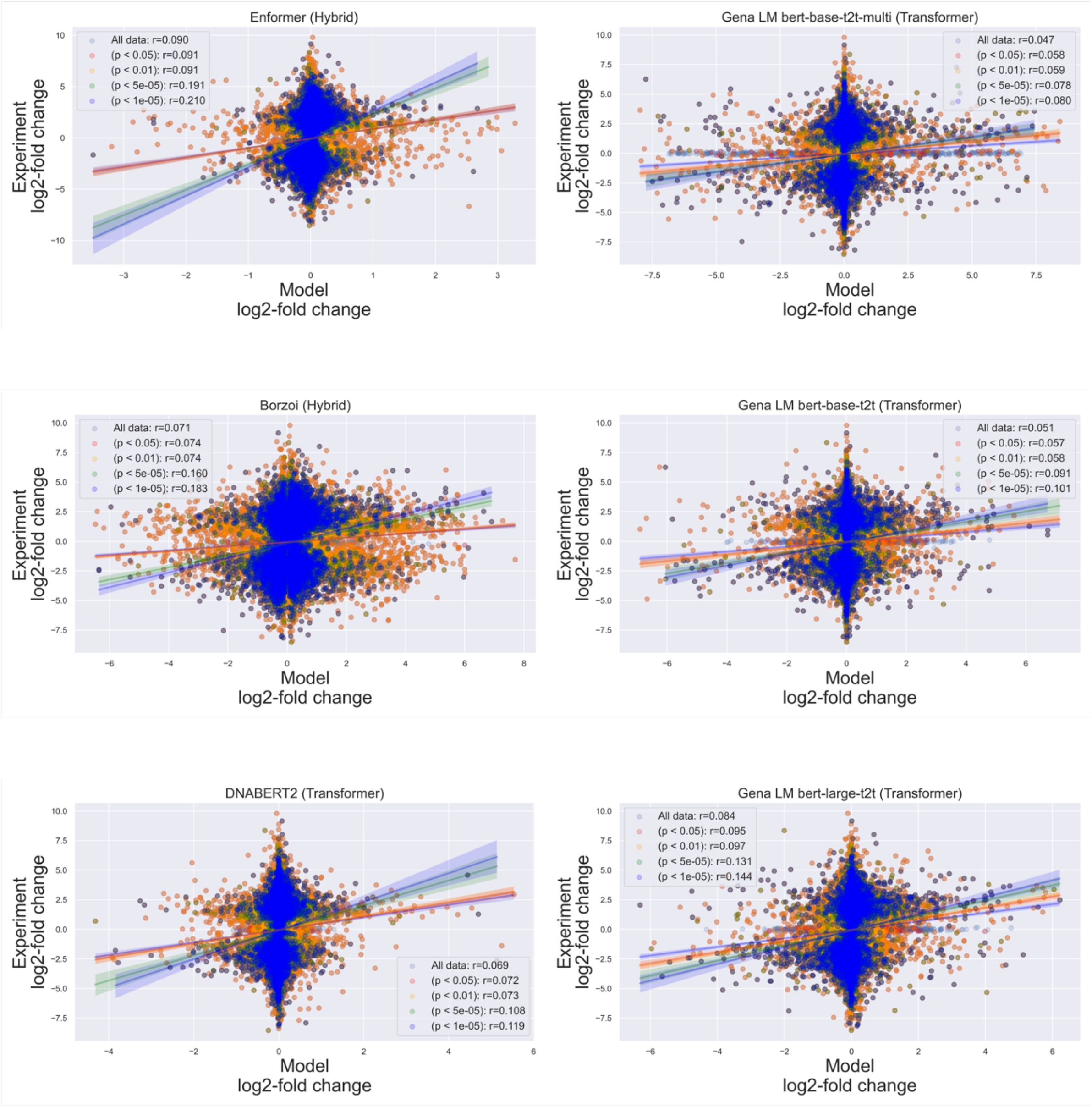


***Figure S3a****: Model performance relative to experimental data significance, using top models from each architecture category.*

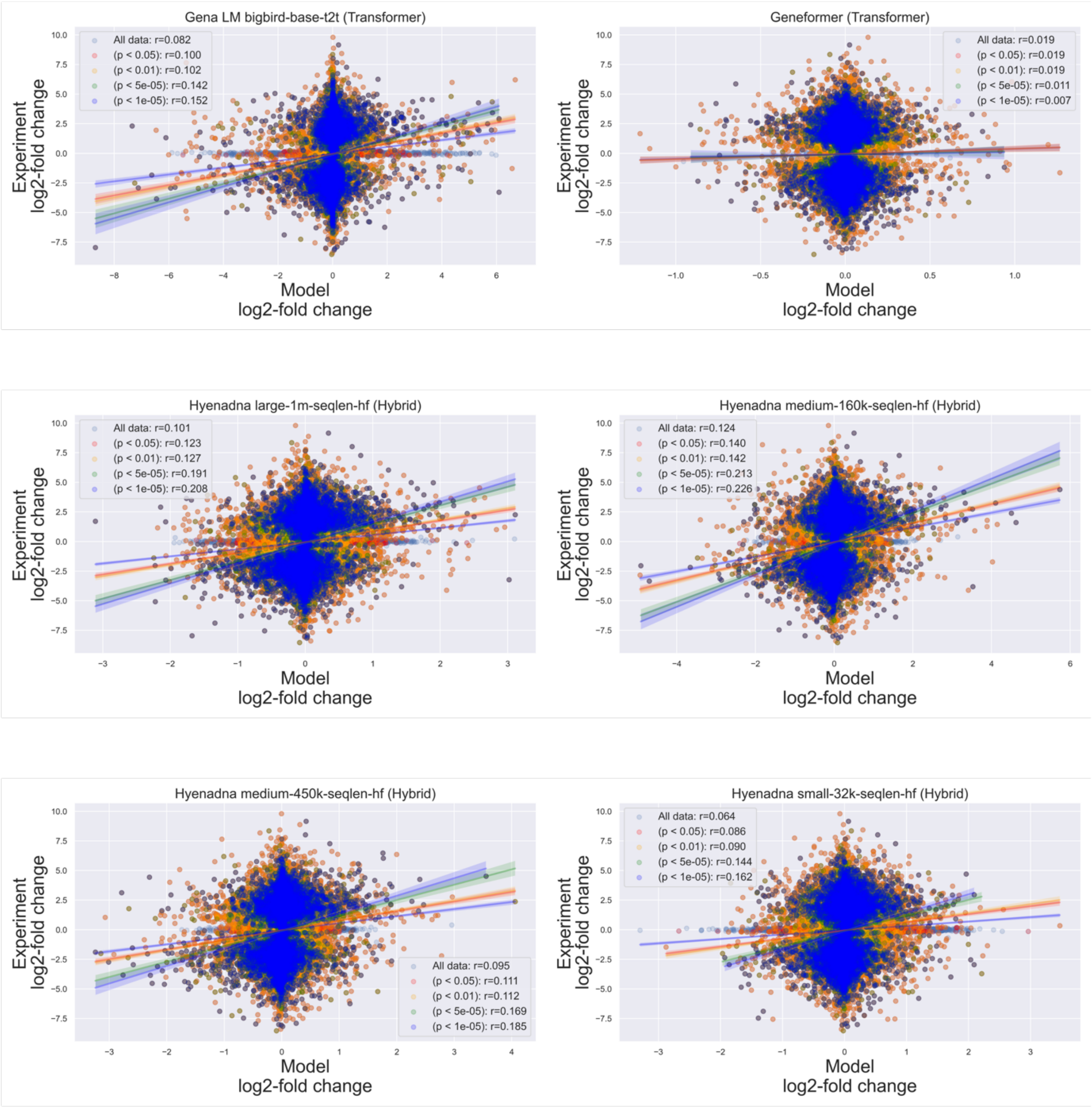


***Figure S3b***: *Model performance relative to experimental data significance, using top models from each architecture category.*

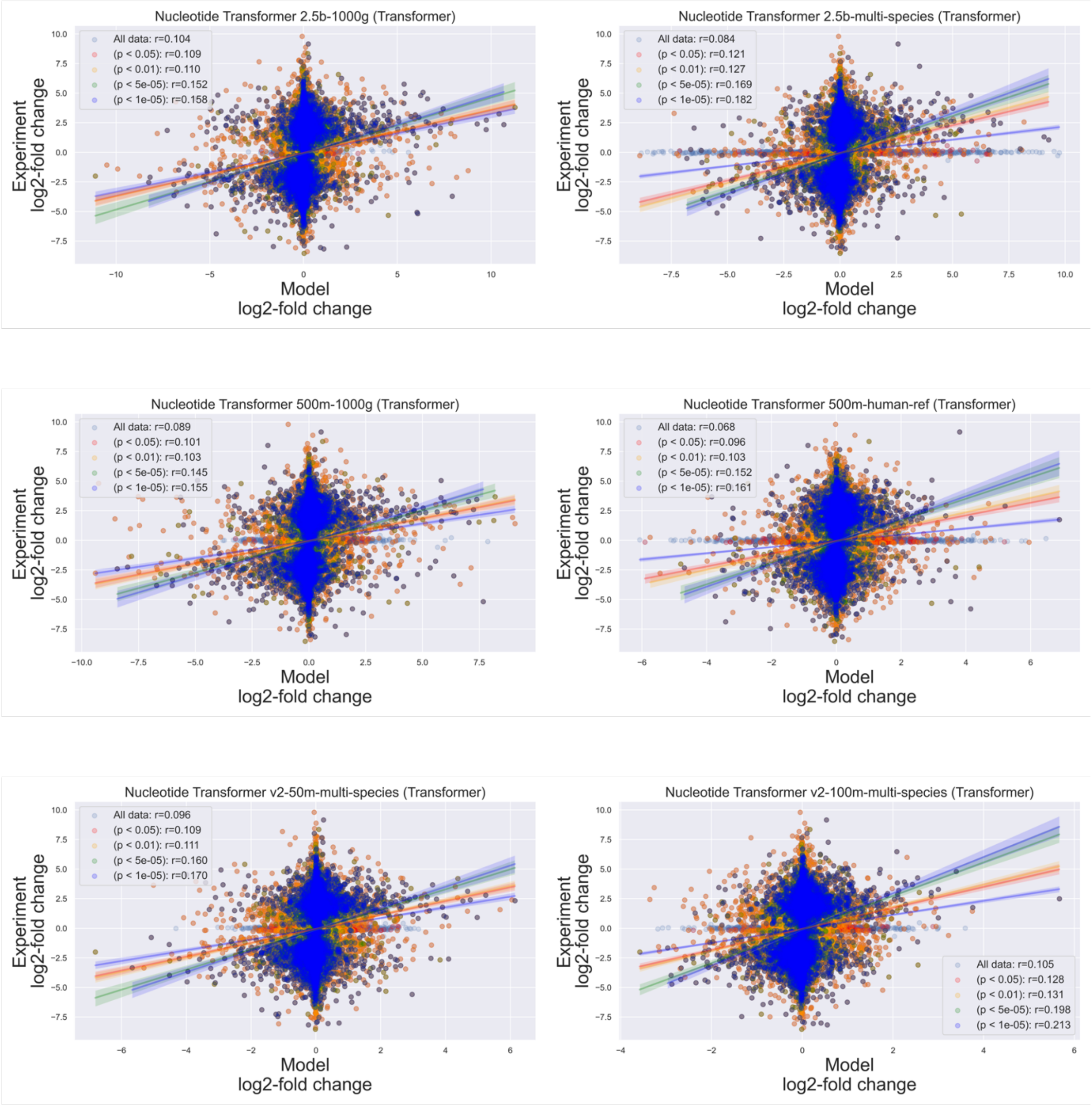


***Figure S3c**: Model performance relative to experimental data significance, using top models from each architecture category.*

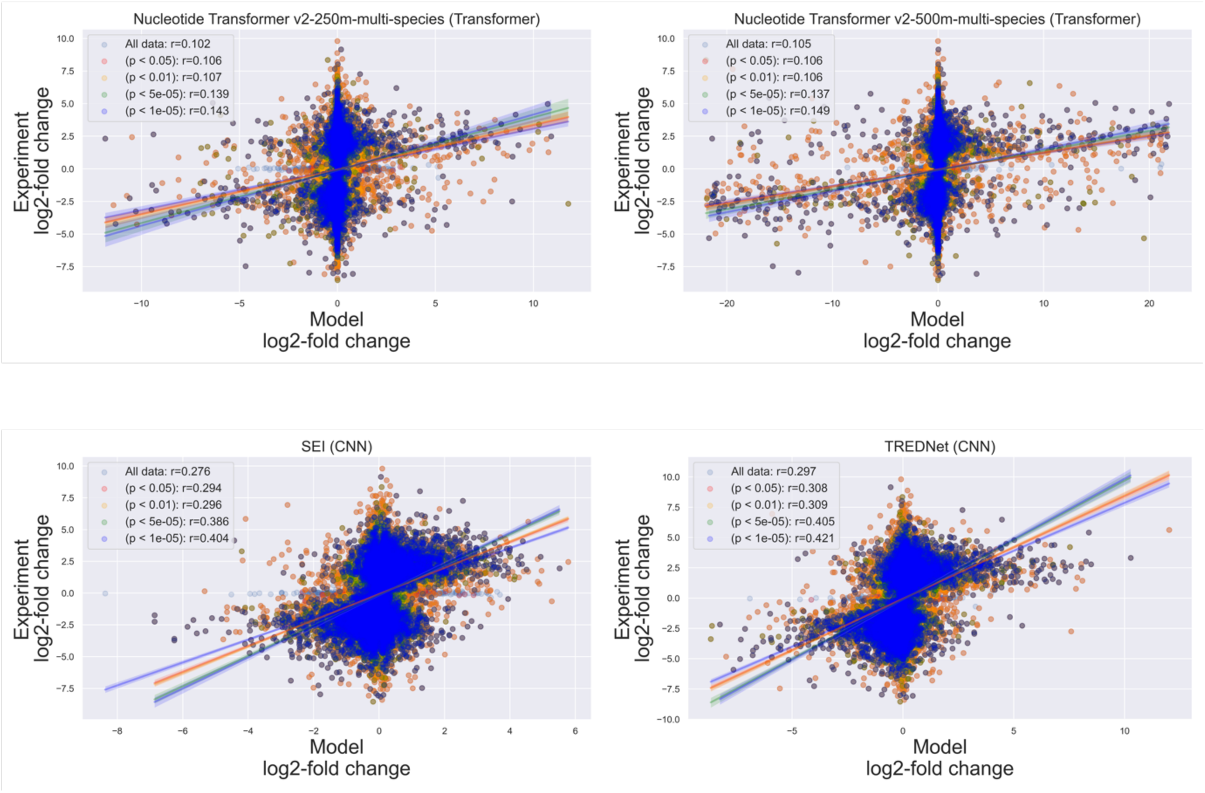


***Figure S3d**: Model performance relative to experimental data significance, using top models from each architecture category.*

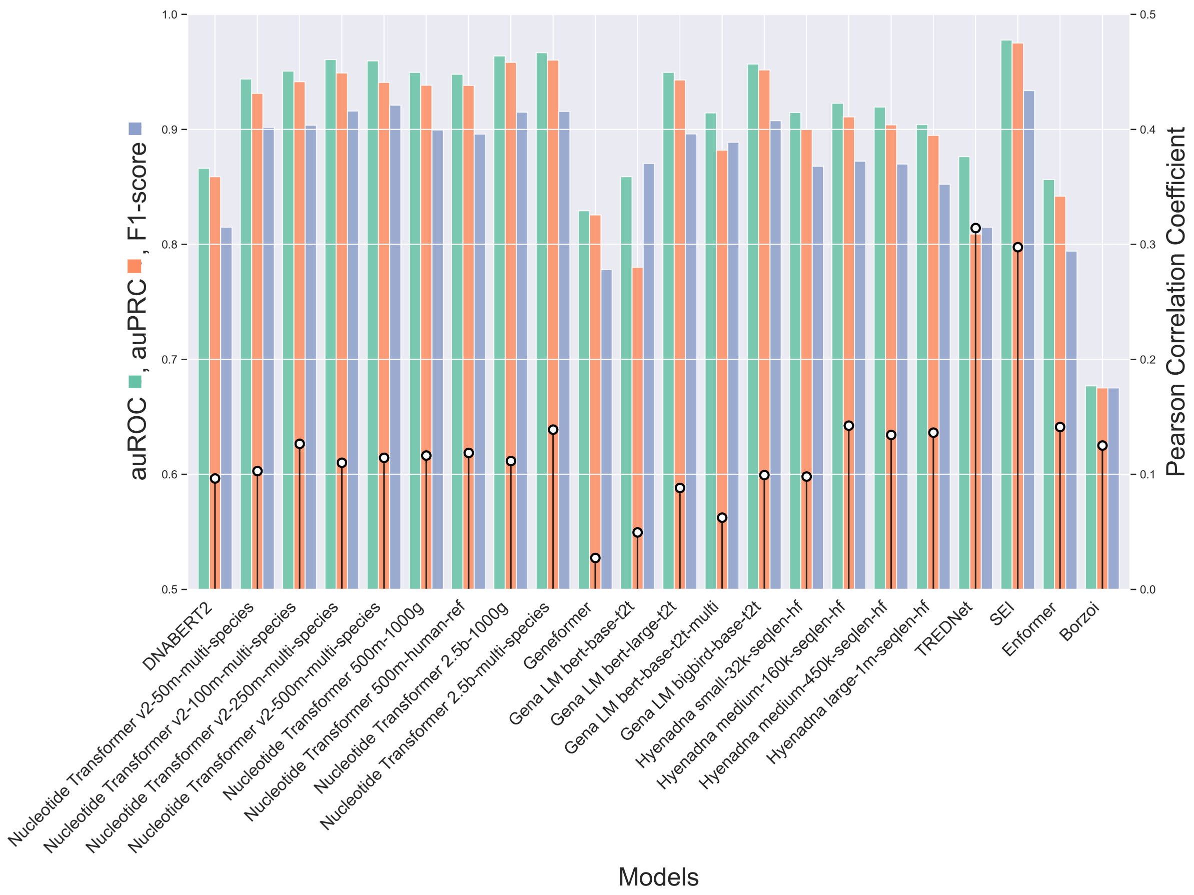


***Figure S4***: *Performance comparison of different models for enhancer detection in HepG2 cell across multiple metrics. Left axis: Area Under the Receiver Operating Characteristic Curve (auROC, green bars), Area Under the Precision-Recall Curve (auPRC, orange bars), and F1-score (blue bars). Right axis: Pearson Correlation Coefficient (black pins).*
